# Large haploblocks underlie rapid, repeatable adaptation during evolutionary rescue

**DOI:** 10.64898/2026.11.15.766882

**Authors:** Brian A. Kissmer, Amy L. Springer, Alia Donley, Zachariah Gompert

## Abstract

Evolutionary rescue can prevent population extinction following abrupt environmental change, but the genetic architectures that enable rapid adaptation under simultaneous strong selection and severe demographic bottlenecks remain poorly resolved. In particular, rescue may rely on either dispersed adaptive variants or clusters of tightly linked alleles that can respond to selection despite strong drift. We investigated the genetic architecture of evolutionary rescue in *Callosobruchus maculatus*, the cowpea seed beetle, following a shift to lentil, a novel and stressful host. To do so, we combined experimental evolution, whole-genome sequencing, backcross introgression, early-generation genomic sampling, association mapping, and transcriptomics. Initial adaptation produced widespread genomic differentiation, but backcrossing localized evidence for adaptive variation primarily to large regions on chromosomes 1 and 9. These regions contained large haploblocks with patterns consistent with suppressed recombination, possibly due to structural variation. On chromosome 9, a rare haplotype increased from approximately 0.5% to 48% in a single generation in one genetic background, consistent with extremely strong selection (*s* ≈ 0.9995). On chromosome 1, a 14 Mb haploblock contained nearly all genetic variation for two fitness-related traits and was associated with differential gene expression. Across genetic backgrounds, chromosome 1 was repeatedly implicated in adaptation, whereas lines from one source population additionally involved chromosome 9. Together, these results show that evolutionary rescue can rely on large, non-recombining haploblocks, providing a mechanism by which clustered genetic architectures can respond rapidly and repeatably despite severe population bottlenecks.

## Main

Rapid adaptation can facilitate population persistence in changing environments, but its genetic architecture varies among systems [1–3]. Empirical studies have documented architectures ranging from many dispersed, small-effect alleles to a few large-effect alleles concentrated within specific genomic regions [4–8]. However, the evolutionary processes that determine where along this continuum adaptive responses fall remain poorly understood [9, 10].

Evolutionary rescue represents an extreme case of rapid adaptation in which adaptation prevents population extinction [11–14]. This process commonly involves strong selection and demographic decline, with the latter imposing severe population bottlenecks and strong genetic drift [15–17]. Theory thus provides specific predictions about the genetic architectures most likely to contribute to adaptation under these conditions. For example, strong drift can overwhelm selection on alleles with small fitness effects [18–20], potentially biasing adaptive responses towards variants experiencing stronger selection [5, 9, 21]. Consequently, when selection on individual alleles is insufficient to overcome strong drift, clusters of adaptive alleles in tight linkage acting as a single evolutionary unit may instead contribute disproportionately to adaptation [9, 22–24]. Thus, evolutionary rescue may be facilitated by “clustered” genetic architectures, in which tightly linked alleles collectively experience selection strong enough to enable an adaptive response despite severe bottleneck-induced drift.

Elucidating the genetic architecture of adaptation during evolutionary rescue remains challenging, as the severe population bottlenecks that often accompany rescue also hamper efforts to resolve its genetic basis [11, 17, 25]. Population decline accelerates non-adaptive allele frequency changes through drift, and can generate extensive genome-wide linkage disequilibrium (LD) [26–28]. This high LD may cause functionally neutral genomic regions to experience selection indirectly due to statistical associations with causal loci (i.e., indirect selection) [26, 29]. Because drift and LD obscure the causes of genetic differentiation during evolutionary rescue, the genetic architectures underlying this process remain largely unresolved.

Experimental evolution provides a framework to study evolutionary rescue during adaptation to controlled, severe, and well-defined selective pressures [30–33]. By exposing replicate lines to these pressures in controlled environments, evolutionary change caused by adaptation can be effectively characterized. Yet even in laboratory conditions, it remains difficult to identify causal genomic regions because bottleneck-induced drift and indirect selection often severely confound signatures of adaptive evolution [34, 35]. To address this, strategic genetic crosses can refine signatures of selection following a strong bottleneck [36, 37]. One such crossing design, involving repeated backcrossing of ancestral lines into adapted lines experiencing ongoing selection in the novel environment, can break down bottleneck-induced LD. This can erode differentiation at regions that evolved primarily due to stochastic processes, thereby refining signatures of adaptive evolution [38]. Integration of experimental evolution and backcrossing approaches remains rare; doing so may provide a powerful approach to resolve the genetic architecture of evolutionary rescue.

The cowpea seed beetle (*Callosobruchus maculatus*) is a tractable model system for the study of evolutionary rescue [39–41]. A cosmopolitan agricultural pest that infests stored grains, *C. maculatus* can adapt to novel hosts of varying suitability [42, 43]. Green lentil (*Lens culinaris*) represents a highly challenging resource with very low initial larval survival (<3%) compared to that on their native cowpea host (*Vigna unguiculata*, >97% survival) [44]. Despite this initial population bottleneck, replicate lines reared on lentil recover fitness approaching that of cowpea ancestors within 20 generations [40]. This represents a case of rapid evolutionary rescue in which genetic adaptation halts and reverses demographic decline [11].

To explore the repeatability of evolutionary rescue across distinct genetic backgrounds, Springer et al. [41] shifted replicate lines of *C. maculatus* originating from Burkina Faso (BF), Brazil (BZ), and California (CA) from cowpea to green lentil. Genetic variants putatively affected by selection were dispersed throughout the genome, yet two chromosomes (1 and 9) harbored very wide peaks of parallel differentiation across many replicate lines, spanning nearly their entire lengths. Importantly, introduction to lentil imposed an extreme bottleneck, reducing census population sizes from approximately 2,000 to as low as ten individuals. Consequently, pervasive drift and indirect selection strongly confounded signatures of F_ST_ and allele frequency change arising directly from adaptation [41]. While genetic variants within the large peaks on chromosomes 1 and 9 likely played a substantial role in lentil adaptation, the origin of the chromosome-wide LD remains unclear. It could reflect selection increasing the frequency of rare haplotypes maintained by suppressed recombination, potentially due to large inversions, or instead result largely from drift during the extreme population bottleneck. Additionally, it is unknown whether smaller, local peaks of differentiation that were detected arose predominantly from drift or selection, leaving the genetic architecture of lentil adaptation unresolved.

To better characterize the genetic architecture of evolutionary rescue during adaptation to green lentil, we first used whole-genome sequencing to capture high-resolution patterns of genetic differentiation in the lentil-adapted lines established in Springer et al. [41] (Figure 1b). We then created backcross introgression lines (BILs) from these lineages to refine and localize the genomic regions associated with adaptation. We combined these analyses with sequence data collected from the earliest stages of adaptation to test for signals consistent with expectations of adaptive structural variation. Finally, we combined genomewide association (GWA) mapping and transcriptomic data to characterize the phenotypic effects of putatively adaptive regions during lentil adaptation (Figure 1). We used these extensive datasets to address three questions regarding the genetic architecture of evolutionary rescue during adaptation to lentil: (i) Are genomic regions that most resist introgression clustered within the broad peaks on chromosomes 1 and 9, or are they dispersed across the genome? (ii) Do patterns of local population genetic structure on chromosomes 1 and 9 support a role for structural variants in addition to bottleneck-generated LD? (iii) Do independent lines of evidence from backcrossing, fitness-associated genetic variation, and gene expression support a clustered architecture of evolutionary rescue?

**Figure 1:**
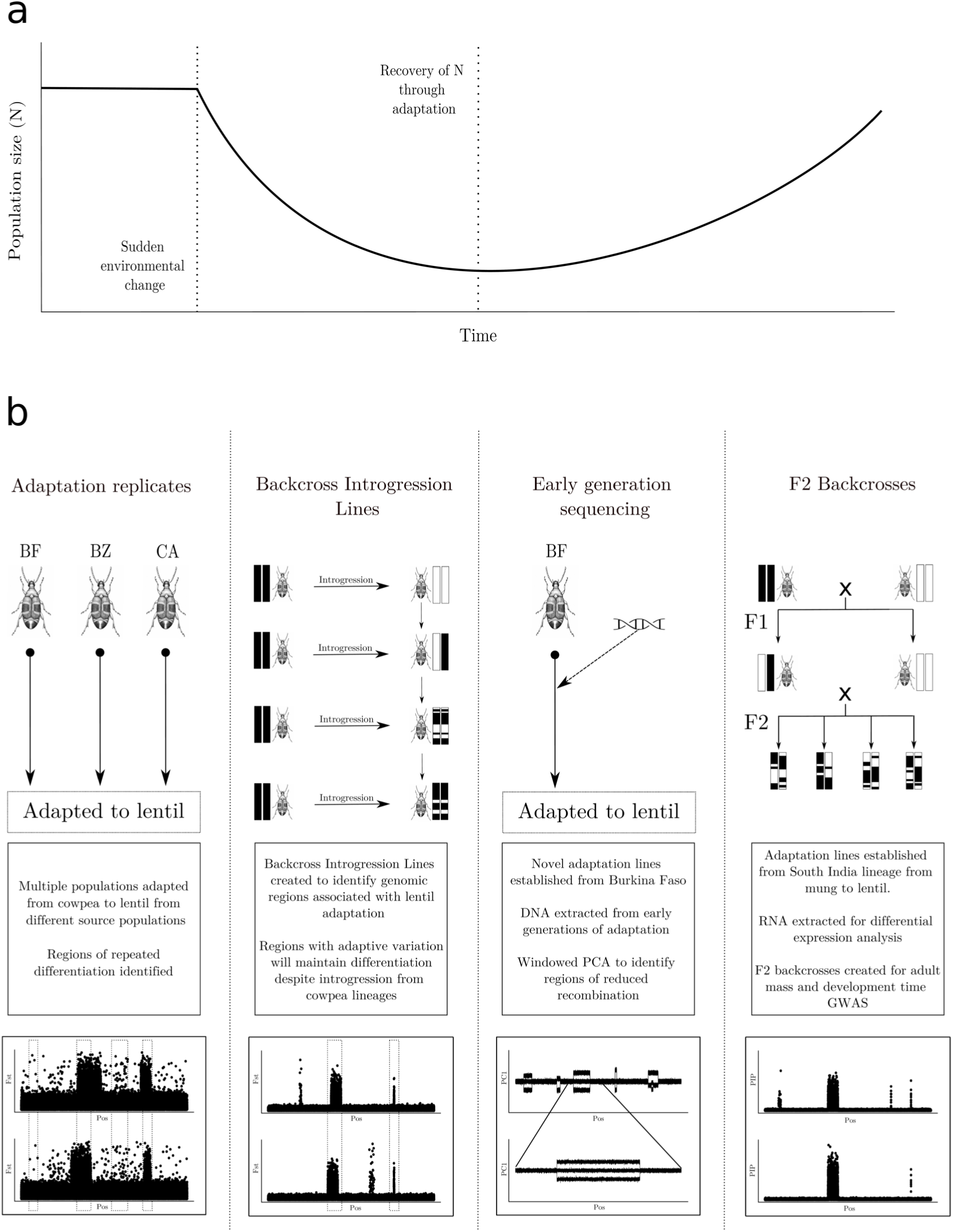
Conceptual framework. a, Stages of evolutionary rescue. Here, population size (*N*) is recovered due to adaptation following demographic decline. b, Experimental design. Replicate seed beetle lines from three geographic origins (BF, BZ, and CA) were shifted to lentil and sequenced with pooled whole-genome sequencing to identify regions of repeated differentiation. Backcross introgression lines (BILs) were then created to identify genomic regions associated with adaptation to lentil. Additional BF lines were shifted to lentil and sequenced upon the initial stages of colonization to identify putatively adaptive haploblocks. Next, differential expression data were evaluated to identify expression outliers associated with survival on lentil. Finally, genome-wide association (GWA) data in backcross mapping lines were used to elucidate the genetic basis of fitness components.

## Results

### Parallel genomic differentiation was widespread following lentil adaptation

To identify repeatedly differentiated genomic regions following lentil adaptation, we resequenced lines reared on lentil for 20 generations, established in Springer et al. [41]– comprising three BF lines, two BZ lentil lines, and two CA lentil lines–along with the cowpea-adapted founders corresponding to each line. The pooled whole genome sequence (WGS) data generated here was used to complement initial genotype-by-sequencing (GBS) data with increased genomic resolution of genetic differentiation. F_ST_ estimated from the 21.4 million SNPs generated from WGS suggests moderate differentiation between the ancestral source lines of BZ and CA from BF (0.164 and 0.156, respectively) and relatively low differentiation between BZ and CA (0.085).

Genome-wide F_ST_ between the founder lines and their lentil-adapted descendants was lower on average than that between the founder populations, ranging from 0.033 (CA replicate 2) to 0.100 (BF replicate 3). Still, the number of SNPs exhibiting greater genetic differentiation than expected by drift alone ranged from 98.9 thousand (CA replicate 2) to 346.9 thousand (BZ replicate 2) after false-discovery rate (FDR) correction (*P <* 0.05). Parallel differentiation across lines was greater than expected by chance in 3,493 of 86,527 10 kb non-overlapping windows (*P <* 0.05, FDR-corrected). Although these windows were dispersed across the genome, 2,428 (69.5%) and 691 (19.8%) were located on chromosomes 1 and 9, respectively.

We next used a hidden Markov model to identify larger contiguous genomic regions of parallel differentiation (RPDs). This analysis identified 24 RPDs ranging from 80 kb to 40.7 Mb in length (Figure 2a). These RPDs were primarily located on chromosomes 1, 5, and 9, and permutation testing confirmed that RPDs on these three chromosomes repeatedly contained higher F_ST_ (estimated in 10 kb windows) than expected by chance compared to regions of similar size (*P <* 0.05) (Table 1). Across replicate lines, the RPDs contained 30% to 88% of the SNPs putatively affected by selection, despite spanning only 15% of the genome. Large RPDs extended from 15.8–44.8 Mb and 54.4–95.1 Mb on chromosome 1, and from 54.5–79.2 Mb on chromosome 9.

**Figure 2:**
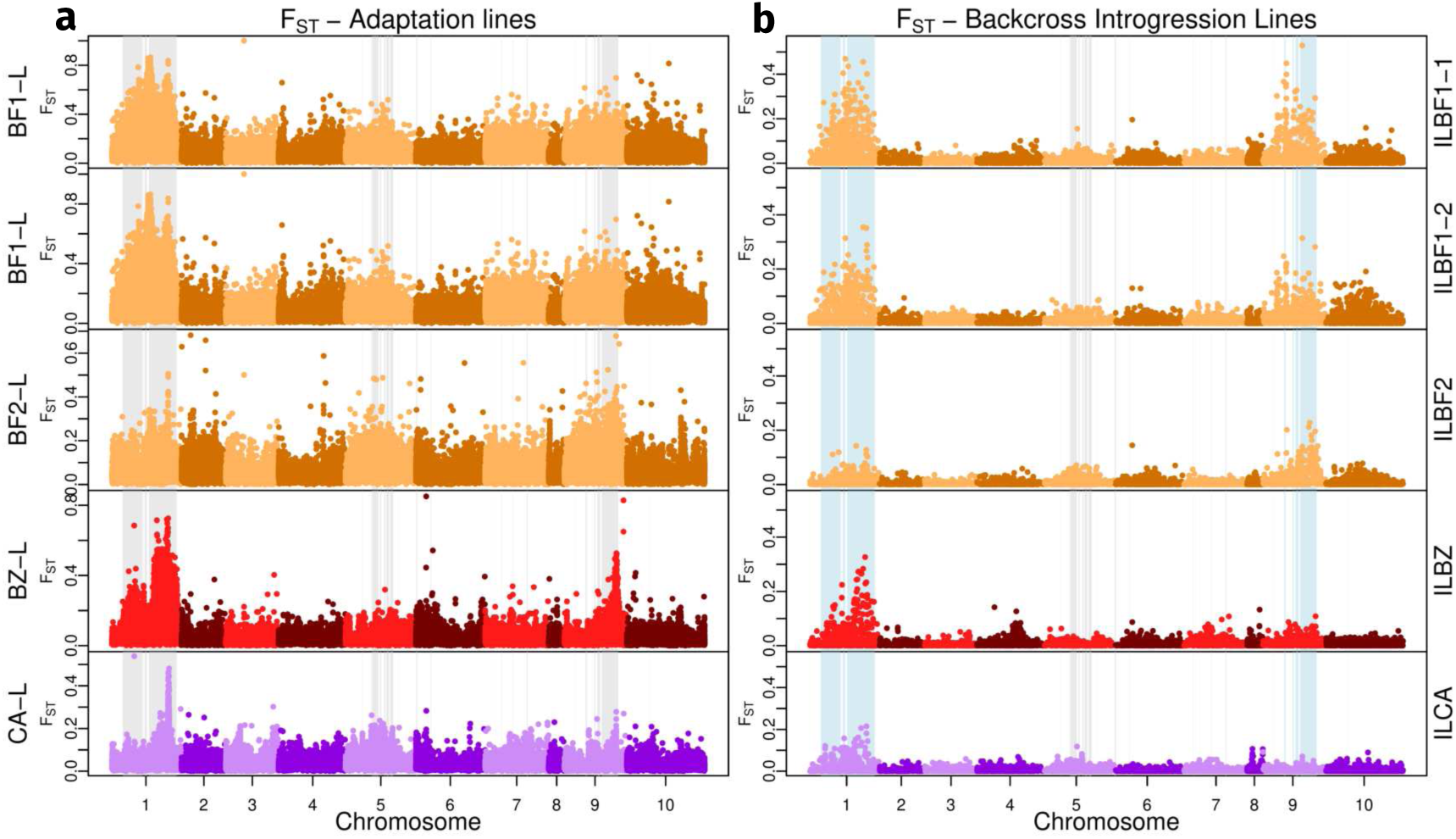
Backcross introgression lines reveal clustered genetic architectures. a, F_ST_ in the pooled WGS lentil-adapted lines used to create the BILs in 10 kb windows. Shading highlights regions that exhibit parallel differentiation across all lines. b, F_ST_ in the BILs. Blue shaded regions maintained differentiation despite introgression in the BILs.

**Table 1:** Regions of parallel differentiation (RPDs) on chromosomes 1 and 9 maintain differentiation despite backcrossing. Summary of the 24 genome-wide RPDs identified from WGS data, alongside significance results following permutation testing in the BILs. For each BIL replicate, RPDs for which *F_st_*was higher than expected by chance are denoted with *α*, and those with reduced effective migration rates are denoted with *β*. Entries for RPDs in which both conditions are true are highlighted in bold.

| **RPD** | **chr** | **Start (Mb)** | **End (Mb)** | **Length (Mb)** | **ILBF1-1** | **ILBF1-2** | **ILBF2** | **ILBZ** | **ILCA** |
|---|---|---|---|---|---|---|---|---|---|
| 1 | 1 | 15.81 | 44.74 | 28.93 | α | α | - | - | - |
| 2 | 1 | 46.26 | 46.87 | 0.61 | α | - | - | - | - |
| 3 | 1 | 48.21 | 51.57 | 3.36 | **α, β** | α | - | α | α |
| 4 | 1 | 54.40 | 95.12 | 40.72 | **α, β** | **α, β** | α | **α, β** | α |
| 5 | 5 | 24.69 | 25.15 | 0.46 | - | - | - | - | - |
| 6 | 5 | 36.92 | 37.56 | 0.64 | - | - | - | - | - |
| 7 | 5 | 38.90 | 48.62 | 9.72 | - | - | α | - | - |
| 8 | 5 | 50.63 | 52.52 | 1.89 | - | - | - | - | - |
| 9 | 5 | 56.55 | 58.45 | 1.90 | - | - | - | - | - |
| 10 | 5 | 59.93 | 63.79 | 3.86 | - | - | - | - | - |
| 11 | 5 | 65.63 | 70.08 | 4.45 | - | - | - | - | - |
| 12 | 5 | 77.23 | 77.44 | 0.21 | - | - | - | - | - |
| 13 | 6 | 0.55 | 2.26 | 1.71 | - | - | - | - | - |
| 14 | 6 | 21.60 | 22.36 | 0.76 | - | - | - | - | - |
| 15 | 7 | 14.67 | 15.02 | 0.35 | - | - | - | - | - |
| 16 | 7 | 23.86 | 24.28 | 0.42 | - | - | - | - | β |
| 17 | 7 | 40.70 | 41.29 | 0.59 | - | - | - | - | - |
| 18 | 7 | 61.38 | 62.72 | 1.34 | - | - | - | - | - |
| 19 | 9 | 14.71 | 14.79 | 0.08 | - | - | - | - | - |
| 20 | 9 | 31.27 | 33.01 | 1.74 | - | - | - | - | - |
| 21 | 9 | 43.89 | 45.57 | 1.68 | α | α | - | - | - |
| 22 | 9 | 48.03 | 52.55 | 4.52 | α | - | α | - | - |
| 23 | 9 | 54.52 | 79.20 | 24.68 | α | - | **α, β** | - | - |
| 24 | 10 | 32.31 | 32.63 | 0.32 | - | - | - | - | β |

### Adaptive variation was repeatedly associated with the same genomic regions

Repeated differentiation suggests a role for nonrandom evolution in the RPDs, yet it is unclear to what extent variation associated with lentil adaptation is localized to these regions. Thus, backcross introgression lines (BILs) were created from the adaptation lines to refine the signatures of selection in the RPDs (Figure 1). This was done by hybridizing the lentil-adapted lineages with their ancestral cowpea lines and then rearing the offspring on lentil, repeating backcrossing to the cowpea lines three times over seven generations. Alternating generations of backcrossing and within-population mating allowed a generation of recombination, potentially reducing the size of ancestral haploblocks—providing recombination suppression (as might occur with structural variation) did not maintain LD. Five BILs were established to assess patterns of evolutionary parallelism using one BZ line (ILBZ), one CA line (ILCA), and two lines of BF (ILBF1 and ILBF2). Two backcross lines were created from the same post-adaptation BF line (ILBF1-1 and ILBF1-2) to test the repeatability of introgression during backcrossing. Beetles from each BIL were sequenced using Genotyping-by-Sequencing (GBS) from which allele frequencies and F_ST_ (in 10 kb windows) were estimated. These data were used to quantify effective migration, that is, the rate of genetic influx into a population following migration, and to assess genetic differentiation between the cowpea parental lines and their respective BILs.

Overall, F_ST_ between cowpea-adapted founders and the BILs was reduced by roughly half through backcrossing relative to that between the founders and the lentil-adapted lines, consistent with the homogenizing effect of gene flow on genetic variation (Table 2). Although RPDs were distributed across six of the ten *C. maculatus* chromosomes, only those on chromosomes 1 and 9 resisted introgression more strongly than the genomic background in the BILs. The genome-wide effective migration rate was lower than the actual migration rate (*m* = 0.5) for all replicates, ranging from 0.15 (ILCA) to 0.19 (ILBF1-2). Further, permutation tests identified significantly reduced effective migration within RPDs primarily on the distal half of chromosome 1 relative to the genomic background (Figure 3). For example, the largest RPD on chromosome 1, RPD:4 (54.4 - 95.1 Mb), resisted introgression more strongly than the genomic background in ILBF1 and ILBZ, and exhibited marginally significant resistance in ILCA (*P* = 0.06). This RPD additionally harbored higher F_ST_ than expected in all five BILs (*P <* 0.05). Similarly, all BILs except ILBF2 exhibited elevated F_ST_ with RPD:3 on chromosome 1 (48.2 - 51.6 Mb). Conversely, chromosome 9 only exhibited significantly greater resistance to introgression in ILBF2, although all ILBF lines showed higher F_ST_ than expected within RPDs on this chromosome (Figure 2b). Unlike the other BILs, ILBF2 only contained a combined signal of significantly reduced migration and high F_ST_ within RPD:23 (chromosome 9, 54.3 - 79.2 Mb) (Table 1). Finally, ILBF1-1 and ILBF1-2 exhibited near identical patterns of comparatively reduced effective migration in the same RPDs, suggesting that variation among BILs more likely reflects differences in the genetic architecture of adaptation among the lentil-adapted lines than stochastic variation in the outcome of the backcrossing process.

**Figure 3:**
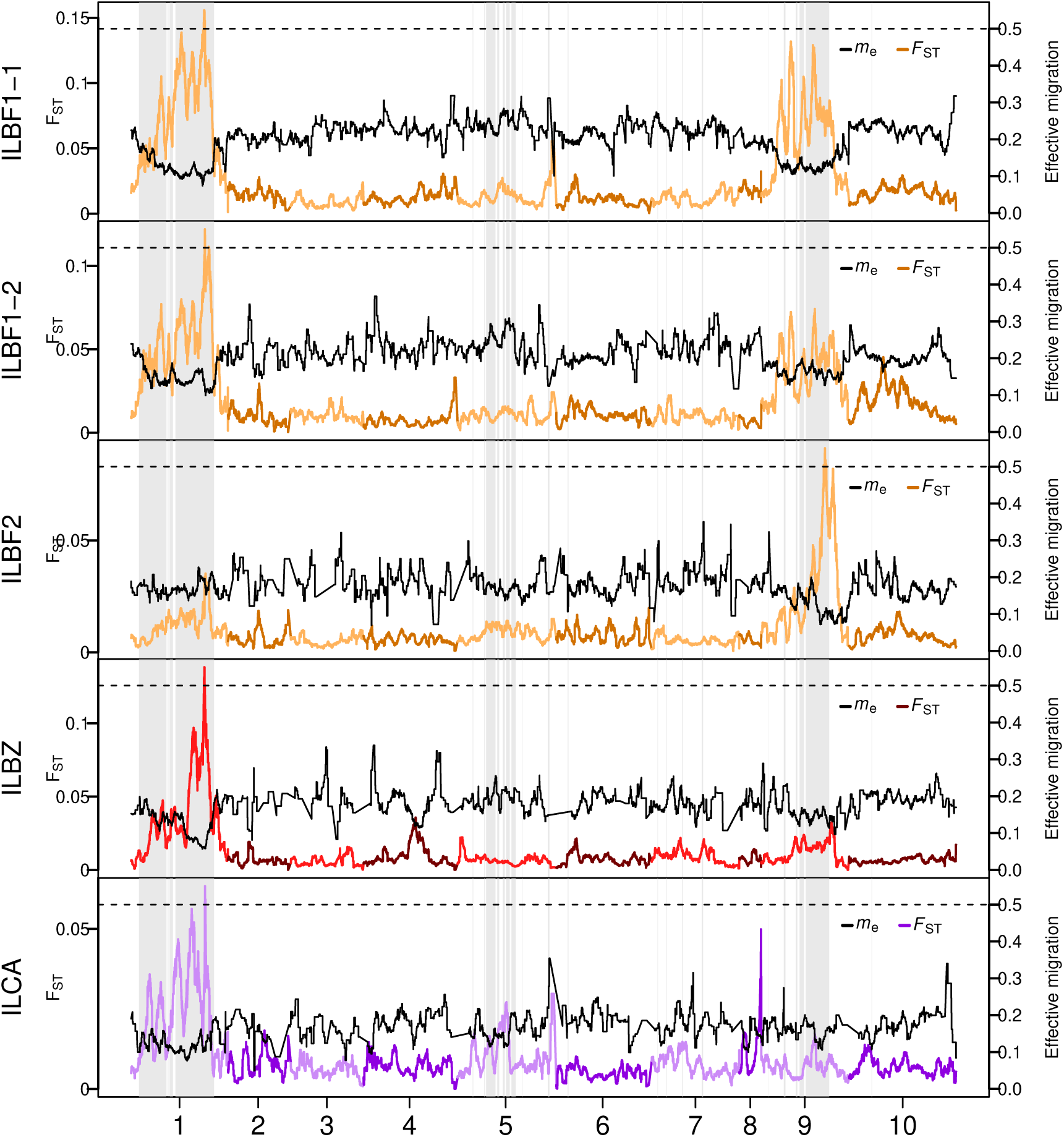
Regions of repeated differentiation resisted introgression relative to the genomic background. Effective migration rate relative to a raw rate of *m* = 0.5 (black) plotted alongside F_ST_ for each BIL, each summarized with 5 Mb overlapping windows in 10 kb steps. Grey bars highlight regions of parallel differentiation (RPDs).

**Table 2:**
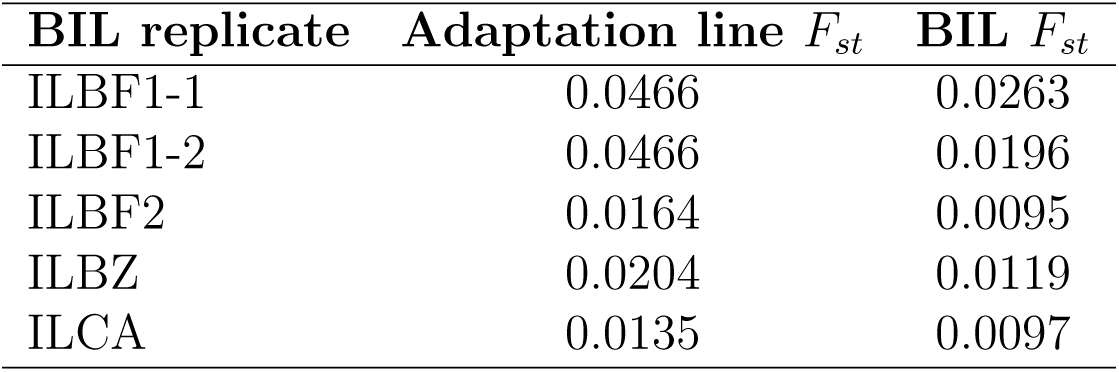
Backcrossing reduces *F_st_*between backcross introgression lines and their cowpea ancestors. *F_st_* between ancestral cowpea-adapted lines and lentil-adapted lines (first column), and between cowpea-adapted lines and backcross introgression lines (second column).

| **BIL replicate** | **Adaptation line** *F<sub>st</sub>* | **BIL** *F<sub>st</sub>* |
|---|---|---|
| ILBF1-1 | 0.0466 | 0.0263 |
| ILBF1-2 | 0.0466 | 0.0196 |
| ILBF2 | 0.0164 | 0.0095 |
| ILBZ | 0.0204 | 0.0119 |
| ILCA | 0.0135 | 0.0097 |

### Regions of reduced recombination underlie adaptive genetic variation

The maintenance of large, introgression-resistant regions on chromosomes 1 and 9 suggested the potential presence of recombination-suppressed haploblocks, as might occur with chromosomal rearrangements. We hypothesized that if such haploblocks drove adaptive differentiation, they would be actively segregating in the first generations of adaptation. To test this, we collected GBS sequence data from the first two generations of surviving beetles following lentil adaptation. The two population replicates used for this, hereafter referred to as BF3 and BF4, originated from a single source BF population, which was chosen for this experiment because BF was the only BIL lineage in which introgression was comparatively reduced (relative to the genomic background) in chromosomes 1 and 9. Using sequence data derived from these early generations, we combined genetic differentiation (F_ST_) and local population genetic structure to identify genomic regions exhibiting patterns consistent with expectations for haploblocks associated with structural variation.

Signatures of genetic differentiation following the host shift were constrained to chromosome 9. Specifically, only the RPDs on chromosome 9 showed signs of differentiation in the first generation (measured as F_ST_) while chromosome 1 was no more differentiated than the genomic background (*P >* 0.05) (Figure 4a). Within the first generation of the BF3 line, local principal component (PC) windows within the peak of differentiation were consistently above the genome-wide 95th quantile for the Calinski-Harabasz index (*k* = 3), a metric that evaluates clustering accuracy, showing strong support for three distinct genetic clusters (Figure 4b). A local PC plot using individuals from both the founder and BF3 F_1_ line confirmed this structure, with three distinct non-overlapping genetic clusters based on SNPS data from 46-62 Mb (Figure 4d), and with similar results in BF4 F_1_ (Figure S1). This 46-62 Mb region additionally possessed higher LD than expected for its size relative to the background of chromosome 9 in both BF3 and BF4 with a value of 0.086 compared to a chromosome-wide background of 0.052 (*P <* 0.05) (Figure 4j). The haploblock coincides approximately with the RPD that resisted introgression in the BF BILs (from 54.5 - 79.2 Mb), suggesting a potential association with lentil adaptation. Taken together, these results are consistent with expectations for large haploblocks associated with structural variants, such as inversions, underlying signals of differentiation in these lines.

**Figure 4:**
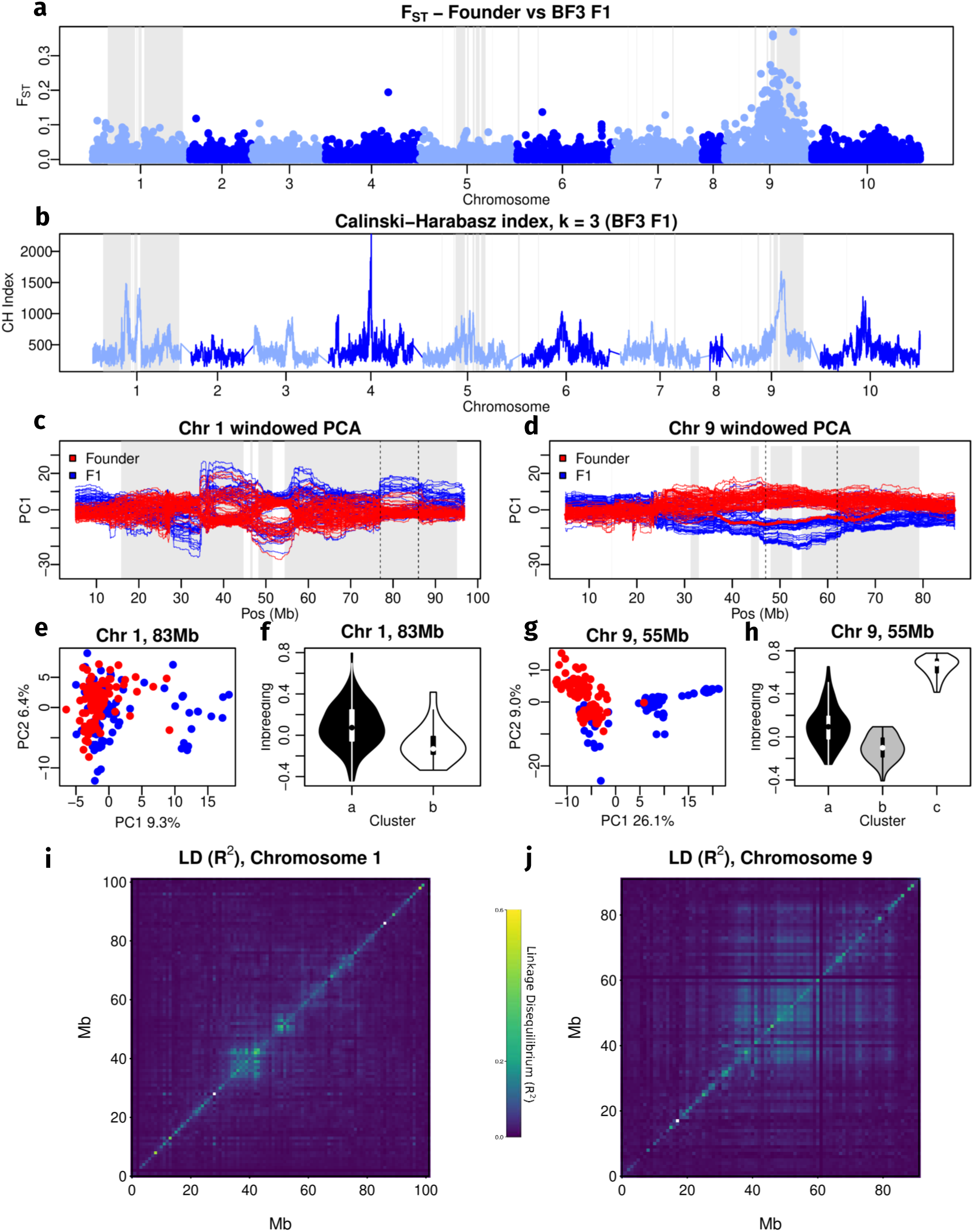
Regions of repeated differentiation resisted introgression during back-crossing. **a**, F_ST_ between the founder line (reared on cowpea) and the BF3 replicate after one generation on green lentil. Grey bars highlight the regions of parallel differentiation (RPDs). b Calinski-Harabasz (CH) index (*k* = 3) estimated for the first component of each 10 Mb local principal component analysis (PCA) window. Windows with larger CH indices fit a model of three clusters better than other windows. c-d, 10 Mb local PCA of chromosomes 1 and 9 for the BF3 replicate. Line color denotes PC1 scores of beetles from F_0_ (red) or F_1_ (blue) generations. A founder individual with the putative heterozygote genotype on chromosome 9 is shown in bold and highlighted with a red arrow (d). e-h, Genotype delineation of large putative haploblocks underlying adaptive regions on chromosomes 1 (e-f) and 9 (g-h). For each, a scatterplot the first two principal components is given at a region of representative structure (e,g) Using genetic clusters identified at these regions, inbreeding coefficients (F_IS_) were estimated within each haploblock (chr1:77-86 Mb and chr9:47-62 Mb respectively) at each location and summarized by cluster in violin plots (f-h). i-j, LD heatmaps (summarized in 1 Mb windows) for chromosomes 1 and 9 in the BF3 F_1_ replicate.

We next tested whether genetic variation for SNPs within this region exhibited patterns consistent with expectations of individuals harboring the homozygous ancestral haploblock genotype, the homozygous lentil-adapted haploblock genotype, or heterozygous genotype. In BF3, the cluster with highest PC1 scores consisted of both founder and F_1_ individuals and had modest inbreeding coefficients (F_IS_) (0.1, s.d. = 0.18), likely representing the reference homozygote haploblock. The middle cluster exhibited the lowest F_IS_ (−0.1, s.d. = 0.12) suggesting a heterozygote genotype, and entirely consisted of F_1_ individuals (except for a single founder individual). The cluster with lowest PC1 values lacked any founder beetles and had the highest F_IS_ (0.65, s.d. = 0.09) indicative of an initially rare, alternate homozygote haploblock (Figure 4g-h). The frequency of each genotype in the founder generation (p/p, p/q, and q/q) was 0.989, 0.011, and 0 (*p* = 0.995, *q* = 0.005), while the frequency of each genotype in the F_1_ generation was 0.29, 0.46, and 0.25 respectively (p = 0.52, q = 0.48). This magnitude of allele frequency shift corresponds to an extreme selection coefficient of *s* ≈ 0.9995, where *s* is defined as the total fitness difference between the alternative homozygotes. This suggests a selection response akin to near lethality of the ancestral allele on lentil. BF4 showed a similar trend in allele frequency change at 0.43 (p/p), 0.43 (p/q), and 0.14 (q/q) after one generation (p = 0.65, q = 0.35). This represents a selection coefficient of *s* ≈ 0.9956, suggesting strong selection against the ancestral haploblock in both lines.

Despite evidence for adaptive variation on chromosome 1 in the BILs, this chromosome did not exhibit elevated F_ST_ in the BF3 and BF4 lines. Additionally, local PC structure and LD did not show patterns consistent with structural variation in RPD:4 (chr1:54-95 Mb). This chromosome further lacked strong evidence of high LD in the RPDs for the early-adaptation lines, despite elevated LD on chromosome 1 in the BILs. It is therefore possible that the signals of differentiation on chromosome 1 in the BILs could be explained by similar haploblocks which were not present in this BF population, suggesting a role of historical contingency in repeatability during adaptation to lentil.

### A large haploblock is responsible for phenotypic variation

We evaluated whether chromosomes 1 and 9 drove phenotypic variation between ancestral and lentil-adapted lines using expression data and genome-wide association mapping (GWA) of backcrossed mapping lines. First, expression data from an additional *C. maculatus* lineage, South India (SI), was reanalyzed following alignment to the updated chromosome-level genome assembly used in this study [45, 46]. RNA expression data were collected from SI lines adapted to lentil for 107 generations. Comparisons were made between mung and lentil adapted lines, reared for a generation on mung to isolate expression differences due to genetics. This revealed that the RPD on chromosome 1 which had resisted introgression relative to the genomic background in the BILs contained more differentially expressed genes than expected by chance (*P <* 0.05, Figure 5a-b). This region contained 13 of the 172 genome-wide significant differentially expressed transcripts with a *P*-value below 0.01 after false-discovery rate correction. Nine of these 13 highly significant loci were upregulated, with an average log fold-change 2.34 (s.d. = 1.78). BLAST results for these genes suggest roles in detoxification, chemoreception, and signaling among other functions, consistent with physiological mechanisms of adaptation associated with novel nutritional regimes (Table S2).

**Figure 5:**
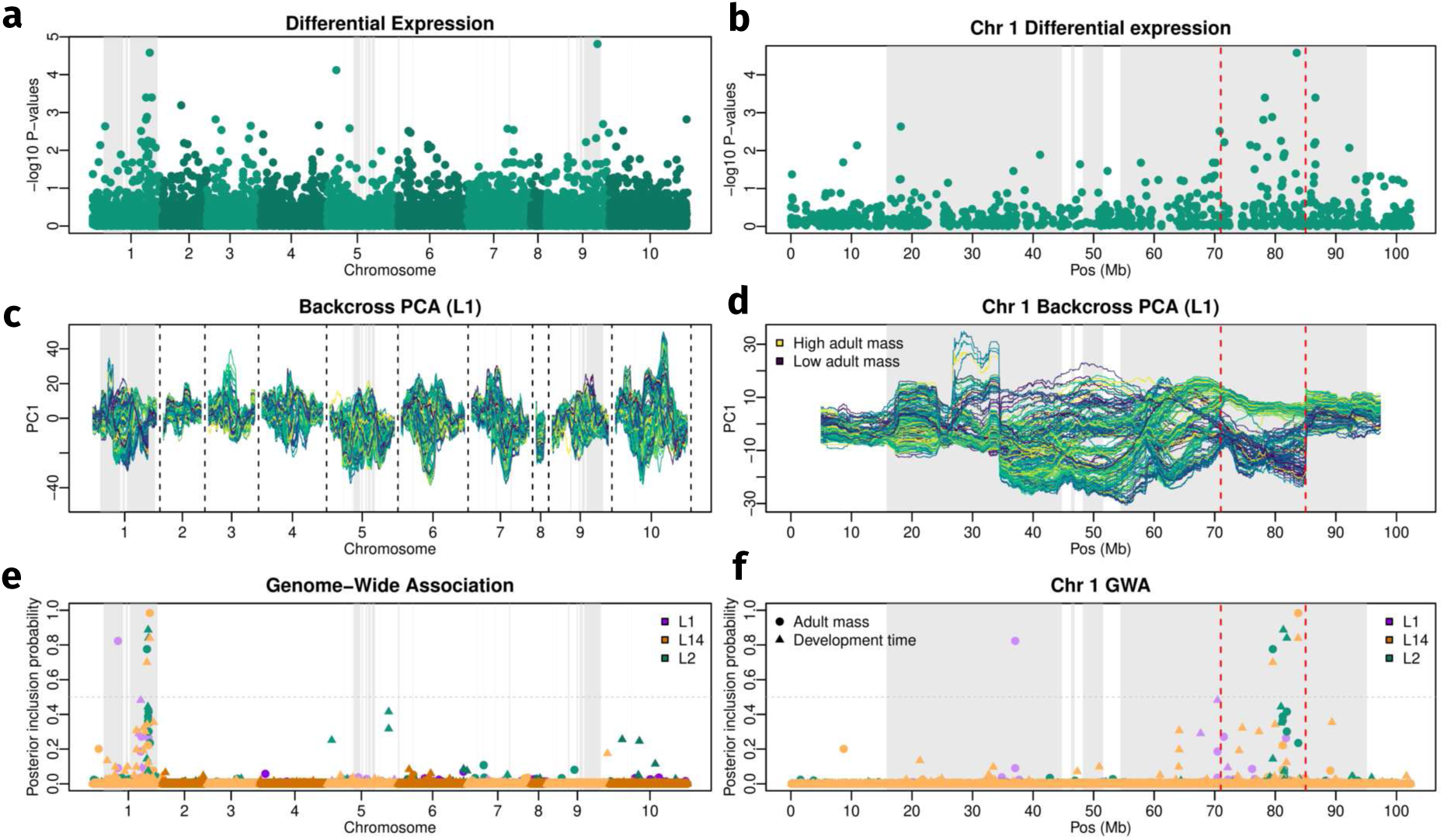
Large haploblock explains phenotypic variation for fitness-relevant traits. Genome-wide (a, c, e) and chromosome 1 (b, d, f) visualization of key characteristics associated with phenotypic differentiation in the SI backcrosses. a-b, Differential expression analysis − log_10_ *P*-values for the SI lineage. c-d, 10 Mb windowed genotype PCA for the L1 backcross replicate. Tracks for PC1 scores are colored based on adult body mass for each individual. e-f, Posterior inclusion probabilities for SNP effects on adult mass (circles) and development time (triangles). Red dotted lines (b, d, f) highlight a large haploblock associated with phenotypic differentiation.

We next used genome-wide association mapping to investigate the genetic basis of fitness components on lentil in three SI mapping populations. Specifically, we used three backcross populations derived from lentil-adapted SI lines that had been maintained on lentil for 146 (L1), 135 (L2), and 38 (L14A) generations. The F_1_ hybrids were generated by crossing each lentil-adapted line with an SI line maintained on its ancestral host, mung bean, and the resulting F_1_ individuals were then backcrossed to the corresponding lentil-adapted line. We mapped development time and adult mass using GBS data and a Bayesian sparse linear mixed model (BSLMM) fit to 27,931 SNPs, allowing us to estimate both polygenic effects and sparse effects attributable to individual loci. Both adult mass and development time showed moderate to high proportions of phenotypic variance explained by additive genetic variation. For adult mass, the proportion of variance explained (PVE), with 95% credible intervals, was 0.51 (0.35 to 0.67), 0.68 (0.55 to 0.81), and 0.82 (0.69 to 0.98) for L1, L2, and L14A, respectively. Development time showed a similar pattern, with PVE estimates of 0.29 (0.17 to 0.43), 0.82 (0.75 to 0.88), and 0.76 (0.69 to 0.82), respectively. A substantial proportion of the additive genetic variance was attributable to loci with measurable individual effects. The proportion of the PVE explained by these sparse effects (PGE), with 95% credible intervals, was 0.68 (0.45 to 0.93) for adult mass and 0.87 (0.58 to 0.99) for development time across all lines. Additive genetic correlations between the two traits were negative in all three lineages: −0.21 (95% c.i. −0.33 to −0.086) for L1, −0.19 (95% c.i. −0.306 to −0.069) for L2, and −0.42 (95% c.i. −0.517 to −0.312) for L14A. The direction of these correlations indicates that alleles associated with shorter development times also tended to be associated with greater adult mass, such that genetic effects on the two traits were aligned in a direction expected to increase fitness on lentil. This pattern is consistent with synergistic pleiotropy, or with linkage among alleles having concordant effects on the two traits. Overall, RPDs on chromosome 1 accounted for much of the mapped genetic variation in these fitness components, supporting an important role for this chromosome in adaptation to lentil.

Both of the fitness components appeared to exhibit clustered genetic architectures, as nearly all (94% - 99.5%) genetic variation contributing to these traits was concentrated on chromosome 1 (Figure 5e-f). For lines L2 and L14A (and for development time in L1), genetic variation associated with the fitness components was entirely localized to the 54-95 Mb RPD. For L1, 80% of genetic variation for adult mass was found in the 16-45 Mb RPD and 17% was contained in the 54-95 Mb RPD. All SNPs for which the posterior inclusion probability (PIP) was > 0.5 (i.e., SNPs with highly credible evidence of measurable effects) were localized between 78-84 Mb, aside from the single SNP with the highest PIP for adult mass in L1, which was located at 37 Mb.

To investigate whether patterns of genetic variation in these SI mapping populations were consistent with the fitness components mapping to recombination-suppressed haploblocks, we quantified local principal component (PC) structure in the backcross lines. As above, we estimated local PC structure using overlapping 10-Mb windows advanced in 10-kb steps. Notably, a large region on chromosome 1 spanning most of the SNPs with high posterior inclusion probabilities (PIPs), with the exception of the adult-mass association in L1 near 37 Mb, showed pronounced local genetic PC clustering between 71 and 85 Mb and elevated LD relative to the remainder of the chromosome (Figure 5c–d). Individuals carrying two copies of this haploblock also exhibited phenotypic differences associated with higher fitness on lentil; on average, they emerged 4.3 days earlier and were 22% more massive than other beetles, with highly significant differences between haploblock homozygotes and other individuals for both traits (*P <* 0.0001). In addition, beetles homozygous for the haploblock exhibited significantly higher F_IS_ than other beetles (0.653 versus −1.21; *P <* 0.0001).

**Figure 6:**
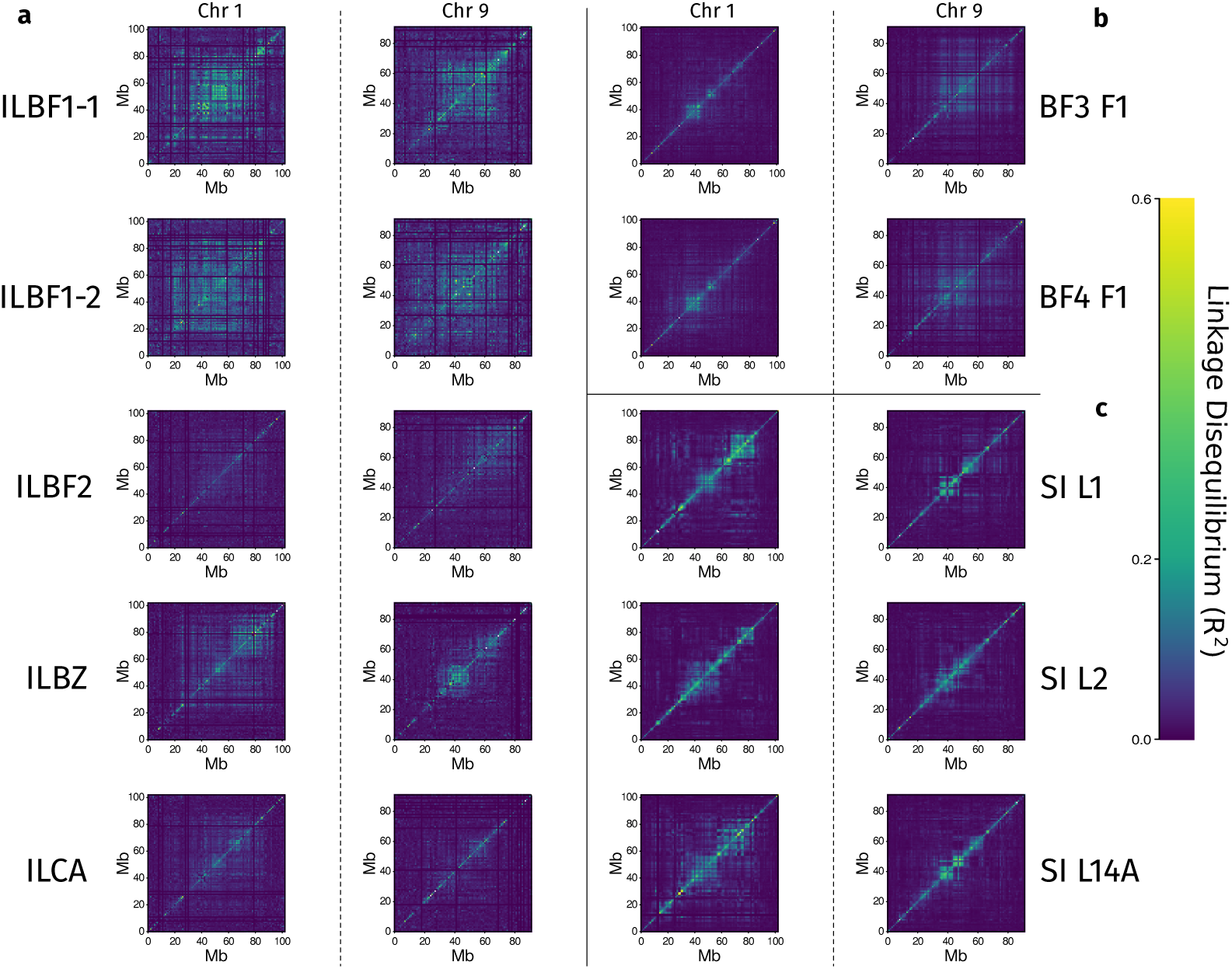
Patterns of linkage disequilibrium (LD) on chromosomes 1 and 9 vary across lineages. LD summarized in 1 Mb blocks for chromosomes 1 and 9 for all GBS-sequenced lines. a, LD for each backcross introgression line. b, LD for each line sequenced at F_1_ following introduction to lentil. c, LD for each SI backcross mapping line.

## Discussion

Here, we used a multifaceted dataset from seed beetle lines adapted to a highly stressful and novel host to elucidate the genetic architecture of evolutionary rescue following a host shift. We showed that genetic architectures were highly clustered and somewhat repeatable, with large haploblocks consistent with expectations of structural variation on chromosomes 1 and 9 underlying adaptation across distantly related lineages. Still, the precise role of these large haploblocks during adaptation to lentil requires further investigation. We discuss hypotheses for the repeated involvement of these two large clusters in light of broader evolutionary theory below.

### Adaptive genetic architecture is clustered and underlaid by large haploblocks

The backcross introgression lines (BILs) revealed that much of the genome-wide F_ST_ present in the initial adaptation lines did not contribute substantially to survival to lentil, as only the RPDs on chromosomes 1 and 9 resisted introgression relative to the genomic background. Large haploblocks with patterns consistent with structural variation were consistently associated with adaptation across datasets [47]. These haploblocks spanned 10–20 Mb, substantially less than the original chromosome-length differentiation peaks, which extended over more than 40 Mb. The broader peaks may therefore reflect both pre-existing LD associated with recombination-suppressed haploblocks and bottleneck-induced indirect selection [29, 48]. The erosion of these peaks following backcrossing was likely caused by introgression from the cowpea-adapted ancestor lines, as recombination broke down associations between the haploblocks and non-adaptive regions. Given that chromosomes 1 and 9 harbored segregating haplotypes during adaptation and explained most of the variation for fitness components, it is unlikely that additional loci dispersed across the genome contributed substantially to evolutionary rescue. These findings align closely with the clustered architecture hypothesis and suggest that large, non-recombining haploblocks (likely associated with structural variation) enabled rapid, repeatable adaptation to lentil despite severe bottlenecks.

The haploblocks associated with evolutionary rescue show behavior consistent with expectations of structural variation. The possible involvement of structural variation here is consistent with a growing body of literature suggesting its potentially central role in adaptation [49–51]. One proposed mechanism by which structural variation facilitates adaptation is through the accumulation of beneficial mutations within non-recombining rearrangements, which can act as co-inherited complexes of advantageous alleles (i.e., supergenes) and facilitate local adaptation despite gene flow [23, 52]. However, the seed beetle lines experienced no gene flow during or leading up to adaptation and encountered a host environment they had rarely, if ever, experienced previously. The haploblocks could have accumulated variants facilitating adaptation to lentil before the four (SI, BF, BZ, and CA) lineages diverged. Yet, this possibility does not explain why alternate haploblocks persisted through prolonged periods on native hosts. Thus, the role of these haploblocks in lentil adaptation likely differs from the conventional supergene model.

The selection coefficient estimated for the haploblock on chromosome 9 during the initial stages of adaptation to lentil suggests exceptionally strong selection (*s* ≈ 0.9995). This greatly exceeds typical selection coefficients observed in empirical studies, which are generally below 0.5 [53]. Given this strength of selection, the maintenance of alternate haploblocks at such low frequencies (*q <* 0.01) for hundreds of generations on their native hosts is surprising. This is particularly striking given evidence of tradeoffs following adaptation to lentil in which fitness on ancestral hosts is reduced [54], which further raises the expectation that the alternate haploblocks should eventually be lost. Balancing selection not tied to host use could nevertheless maintain the polymorphism through several mechanisms. For example, sexual antagonism could maintain the polymorphism [55–57], given its well-characterized role in *C. maculatus* [58–60]. Specifically, if alternative haploblock alleles confer opposing fitness effects in males and females, the polymorphism can be maintained through balancing selection. When haploblocks persist over long periods of time under such conditions, associative overdominance can additionally contribute to long-term maintainence. This is because deleterious recessive alleles can accumulate within haploblocks due to recombination suppression, generating apparent heterozygote advantage and possibly promoting the maintenance of polymorphic haploblocks [61, 62]. Further work is needed to determine how the polymorphism is maintained despite the exceptionally strong selection observed during adaptation to lentil.

### Adaptation to lentil is mostly repeatable

Adaptation exhibited repeated associations with the same genomic regions, involving two large haploblocks on chromosomes 1 and 9. However, differences existed in the extent to which either chromosome was associated with adaptation across source populations and individual lines. Despite extreme (>97%) initial mortality upon the host shift across studies [40, 41, 63], our results implicate genomic background, rather than drift alone, in non-parallel evolutionary outcomes. All lineages except BF exhibited highly repeatable adaptation, consistently involving alleles present on chromosome 1. Conversely, adaptation in BF relied on both the haploblocks on chromosomes 1 and 9, although the extent to which either chromosome was associated with adaptation varied between lines. This work contributes to an increasing body of literature suggesting shared genomic background as a major driver of repeatable evolution [64–66].

Given the rarity of the lentil-adapted haplotypes observed in cowpea-adapted lines, we could not effectively determine whether the alternative haploblock on chromosome 9 initially existed at low frequency in SI, BZ, or CA. If both haploblocks were indeed present, a combination of genetic redundancy and global epistasis could have driven divergent evolutionary outcomes between BF and other lineages [67, 68]. Here, the alternate haploblock on chromosome 9 may exist but have a reduced fitness effect on green lentil in the genomic backgrounds of SI, BZ, or CA. Additionally, this haploblock may not contribute substantially to fitness on lentil given the existence of adaptive variation on chromosome 1 in these lines. Further work is needed to validate the presence of this haploblock in the cowpea-adapted lineages to determine whether genomic background regulates parallel evolution through shared adaptive alleles alone, or alongside complex epistatic interactions with putatively adaptive loci.

## Conclusions

Using an experimentally induced host shift among replicate lines of a cowpea seed beetle, we showed that evolutionary rescue relied on large, non-recombining haploblocks. These haploblocks exhibited patterns consistent with structural variation, existed as low-frequency standing variation prior to adaptation, and controlled nearly all additive variation for two fitness components. Adaptation was largely repeatable despite extreme bottlenecks, with two genomic regions consistently harboring causal variation. Even when the same haploblocks were not repeatedly implicated across lines, adaptation still relied on concentrated genetic architectures. Strong selection on haploblocks with suppressed recombination likely allowed these regions to overcome the high level of bottleneck-induced drift, and repeatable evolution was driven largely by shared genetic variation among lines. This work suggests that structural variation persists over long periods of time and likely facilitates rapid, repeatable evolution during evolutionary rescue on ecological timescales.

## Materials and Methods

### Study system

We used repeated host shifts in the cowpea seed beetle (*Callosobruchus maculatus*, family: Chrysomelidae) to study the genetic basis of evolutionary rescue during rapid adaptation to green lentil. *Callosobruchus maculatus* is a cosmopolitan agricultural pest that originated in Africa and expanded globally due to transportation of infested beans throughout the Holocene [42, 69]. This system is especially well suited for experimental evolution studies as the laboratory environment closely resembles that of their natural environment (i.e., grain legume stores) [70]. While the main host of this species is cowpea (*Vigna unguiculata*), *C. maculatus* has been shown to adapt readily to novel hosts in both agriculture and the laboratory environment [39, 71]. Initial survival on novel hosts is variable, with hosts ranging from suitable to highly challenging [39]. Green lentil (*Lens culinaris*) represents a highly challenging host, very rarely used in natural populations, with initial larval survival ranging from 1-3% [44, 72]. However, laboratory lines reared on lentil have successfully recovered fitness (by proxy of adult mass and development time) relative to their cowpea counterparts in 20 generations or less [44]. Although previous studies have investigated the genetic basis of lentil adaptation in *C. maculatus* [54], including more recent work taking advantage of a recently published chromosome-level reference genome [41, 46], the genomic basis of evolutionary rescue remains incompletely understood.

Our collection consists of populations isolated from Burkina Faso (BF), Brazil (BZ), California (CA), and South India (SI). BF, BZ, and CA were originally collected on cowpea while SI was collected on mung bean (*Vigna radiata*), and each population is maintained on their respective native host. The BF lineage was collected from Ouagadougou, Burkina Faso in 1989 [39, 73]; BZ was collected from Campinas, Brazil in 1975 [74]; CA was collected and maintained by Peter Credland; SI was collected from Tirunelveli, India [75, 76]. Each line was reared in laboratory conditions for well over 300 generations prior to the start of this experiment, so laboratory adaptation is unlikely to confound signatures of adaptation to lentil. The diverse origins of our collections provide an opportunity to test evolutionary repeatability across different genomic backgrounds.

Stock colonies were maintained in 2L Mason jars. Approximately 2,000 adult beetles (measured by volume in an aspirator) were sieved from jars upon emergence and placed in a new jar with approximately 2,000 beans. This allowed for discrete generations, as individual beetles tend to emerge at similar rates in stable conditions (between 24 and 28 days when reared on cowpea). Lines were maintained at 27°C and a 14/10 day cycle in growth chambers in a Percival incubator (model I-36VL). Fluctuations in humidity were caused by a combination of the external environment and the generation of metabolic water by beetle larvae, so dehumidifiers were installed to maintain humidity between 40 and 60% [77].

Previous work on this system has involved numerous host-shift experiments to study adaptation to novel nutritional sources [44, 75, 78]. Although some of this work collected sequence data, genomic analyses were initially constrained by reference genomes that lacked chromosome-level resolution. More recent work has leveraged the chromosome-level reference genome to investigate adaptation in this system [41]. Alongside creating novel experimental lines, this study revisits prior data (as detailed below) by realigning sequence data to the chromosome-level reference genome to further characterize the basis of adaptation to lentil.

### Adaptation experiment

The lentil-adapted lines used in this experiment were established in Springer et al. [41]. To briefly summarize the methods from this study, eleven lentil-adapted replicate lines from each of the three geographic origins (BZ, BF and CA) were established each by placing ∼ 2, 000 beetles from cowpea onto ∼ 2, 000 green lentils in 2L Mason jars. Previously published standard culture procedures were then followed [39], in which offspring were collected upon emergence and placed into a clean jar with another fresh 2,000 lentils. Experimental lines were maintained similarly to their ancestral cowpea lines albeit with adjustments for longer generation times. DNA was extracted from 19-20 beetles per line after fitness had nearly recovered to that of the original cowpea lines (after 20 generations of adaptation), using females to ensure consistent coverage of the X chromosome. DNA was extracted using the Qiagen DNeasy 96 Blood and Tissue Kit according to the manufacturer’s protocol, as detailed in Springer et al. [41].

Of these lines, the current study pooled extracted DNA from three BF lentil lines, two BZ lines, and two CA lines (alongside their ancestral cowpea counterparts) for whole-genome sequencing (WGS), using 48 beetles per pool. Here, DNA concentrations for each sample were measured using the Agilent Biotek system and concentrations were normalized within lines to ensure proportional contributions of DNA sequences from each sample to a population pool. Samples were then sent to the Beijing Genomics Institute (BGI, Beijing, China) for sequencing on the DNBseq platform. This returned 3.89 billion 150 bp reads representing 1.116 trillion sequenced base pairs. Genome coverage for lines ranged from from 104× to 118×.

Raw reads with a Phred score above 33 were retained, and adapter sequences were removed. Whole-genome DNA sequences were aligned to the CASHZR02 reference genome (NCBI accession CASHZR020000000) [46] using bwa (v. 2.0pre2) with the mem2 algorithm under default parameters [79, 80]. Alignments were retained if they possessed a minimum exact seed length of 19 bp and an alignment score greater than 30. The resulting BAM files were sorted and indexed using samtools (version 1.16) [81]. To reduce false-positive variant calling due to insertion-deletion (indel) regions, local realignment around indels was performed using GATK’s RealignerTargetCreator and IndelRealigner (version 3.8) [82]. Alignments for which mapping quality was greater than 20 and individual bases with a quality score above 30 were retained for variant calling using bcftools mpileup (version 1.23-3-g34a49760) [83]. Variant calling was executed using the bcftools call function under the consensus model, with variants being retained if the probability of the homozygous reference genotype was less than 0.01. Custom perl scripts were then used to remove variants with coverage below 500 (∼50× coverage per pool), a mapping quality below 30, and unsigned Mann-Whitney U statistic below 4 for base quality, mapping quality, and read position biases. Finally, triallelic sites were removed. This left 21.4 million SNPs for downstream analyses, from which allele frequencies were calculated using allelic depth.

Allele frequencies from pairs of pre- and post-adapted lineages were used to next determine whether the magnitude of allele frequency change for each SNP deviated from null expectations under a model of pure drift. To do this, varne was used to estimate the contemporary variance effective population size (*N_e_*) using 500 bootstrap replicates [84]. This approach assumes that loci evolved solely in response to genetic drift, so SNPs above the 95th quantile for allele frequency change following adaptation were removed for this analysis. *N_e_* estimates were used to parameterize a null model of evolutionary change expected by drift via a beta-distribution approximation to a Wright-Fisher model [40, 85, 86]. At a given locus, probability of an allele frequency *p_t_* at time *t* is approximated by a beta distribution in which *α* = *p*_0_ ∗ (1 − *F*)*/F* and *β* = (1 − *p*_0_) ∗ (1 − *F*)*/F*, where 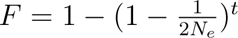. Expectations for the magnitude of change as a result of genetic drift are therefore determined by *N_e_*, time, and initial allele frequency. Thus, our model was parameterized using the number of elapsed generations (*t*), varne estimates of *N_e_*, and allele frequencies measured before and after adaptation. Alleles with a minor allele frequency (MAF) of < 0.01 were removed, and one-tailed probabilities estimated from the beta probability distribution function (pbeta in R) were converted to two-tailed *P*-values using the equation min[*P*_beta_ ∗ 2, (1 − *P*_beta_) ∗ 2] [87]. These *P*-values are interpreted as the chance that allele frequency change at a locus was due to genetic drift alone. Deviations from this null are suggestive of allele frequency change influenced by selection, either directly or indirectly (via linkage disequilibrium) rather than solely by drift.

We next conducted a series of analyses to identify regions of the genome that experienced repeated differentiation across lentil adaptation lines. For this we used the PicMin package in R (version 0.0.0.9000), which tests for repeated evolution against a null expectation of idiosyncratic allele frequency change [88]. PicMin works by identifying windows of SNPs which repeatedly fall into the tails of an empirical *P*-value distribution among a set of populations. Rather than running the model on the original *P*-values generated in the above null model, a 10 kb sliding window-based approach was used to reduce noise and amplify signals which were consistent among contiguous SNPs. To identify windows in which selection likely played a role in differentiation, the number of SNPs with a *P*-value below 0.01 was counted for each population pair. The *log* probability of obtaining the observed number of significant *P*-values given the total number of SNPs in each window was estimated using the cumulative density function of the binomial distribution (pbinom in R). Here, the probability of success was defined as the chance that individual SNPs were significant, equal to the *P*-value of 0.01. These window-based *P*-values were converted to an empirical (rank order) *P*-value distribution. We then used the GenerateNullData function from PicMin with 10,000 replicate draws, using *a* = 0.3 and *b* = 0.5, to obtain the null correlation matrix. The PicMin function was applied to these empirical *P*-values to obtain parametric *P*-values representative of a null model for idiosyncratic evolution, and a false discovery rate (FDR) correction was applied to these final *P*-values [89]. We interpret these final *P*-values as the deviation from the null hypothesis that the observed degree of repeated differentiation across lentil-adapted lines is no greater than expected by chance.

To generate explicit boundaries for regions that experienced parallel genomic differentiation, a hidden Markov model (HMM) was fitted to the negative *log*_10_ *P*-values generated from PicMin using the HiddenMarkov package in R (version 1.8.14) [90]. The HMM was trained to classify windows into one of two states, and a modified version of the Mstep.norm function was used to provide fixed values for means and standard deviations of the two states. For both states the s.d. was set to half the s.d. of the original dataset, while the means for the two states were set to the 5th and 95th percentile of negative *log*_10_ *P*-value distribution, respectively, to distinguish the genomic background from regions exhibiting unusually strong evidence of repeated differentiation. Genomic regions classified into the ‘repeatedly differentiated’ state were combined with neighboring regions within 1 Mb under the assumption that these windows likely diverged together via linkage with the same causal loci, after which 24 regions of parallel differentiation (RPDs) were identified. Finally, a permutation test was conducted to verify whether these regions had higher F_ST_ than the genomic background across lines. Here, null distributions for each RPD were created by counting the number of loci above the 95th percentile of F_ST_ for windows the same size of each RPD across the genome. Sliding windows were used instead of random sampling of genomic loci to preserve the spatial genomic structure of existing F_ST_ outliers.

### Backcross Introgression Lines

We were next interested in the extent to which the regions of parallel differentiation (RPDs) contained genetic variation associated with lentil adaptation. To determine this, backcross introgression lines (BILs) were created from the adaptation lines to identify regions of the genome that harbored genetic variation associated with survival on lentil. Approximately 250 males and females (500 beetles total) from each lentil adapted line were mated with the corresponding number of beetles from their ancestral cowpea line and placed on lentil to lay eggs. This was accomplished by placing individual beans with near-emerged beetles in individual gelatin pill capsules. Pill capsules were scanned for virgin beetles as they emerged, which were separated based on sex. This was done to ensure that the resulting offspring were true hybrids such that the resulting migration rate was as close to 0.5 as possible. Backcrossing occurred at generations 1, 2, and 5 of the BILs. Alternating between generations of backcrossing and random within-population mating allowed for recombination to reduce the size of ancestral haploblocks between generations of backcrossing, provided that recombination was not suppressed (i.e., due to structural variation). After 7 generations, adult females were isolated from the BILs as well as their cowpea and lentil parental lines for genotyping-by-sequencing (GBS).

Genetic differentiation among the parental lines and the BILs was determined using GBS. For this, DNA was extracted from 35 beetles from each BIL and 25 from each parental line using the Qiagen DNeasy 96 kit according to manufacturer’s instructions. While the Y chromosome of *C. maculatus* has been shown to harbor potentially relevant loci [91], females were used for sequencing to guarantee even coverage on the X chromosome. Extracted samples were digested using the EcoRI and MseI restriction digest enzymes then barcoded using the EcoRI library, a collection of 384 unique 8-10 bp internal barcode sequences. PCR was used to amplify restriction fragment libraries, after which a BluePippin (Sage Science, Beverly, MA, USA) was used to isolate fragments between 250-350bp in length ensuring consistent sites were sequenced across the genome for all lineages (Sage Science, USA). The DNA library was sent to the Utah State University Genomics Core facility for sequencing on a NextSeq 2000, which returned 840,833,701 150 bp single-end reads corresponding to approximately 126.12 billion sequenced base pairs (Illumina, USA).

PhiX reads were removed prior to downstream analysis. Reads were aligned to the PhiX genome (NCBI accession NC 001422) using the bwa aln algorithm (version 0.7.17) [79]. Here, the seed length (-l) was set to 20, the maximum mismatches allowed in the seed sequence (-k) was set to 2, and the number of mismatches allowed in the entire sequence (-n) was set to 5. After removal of PhiX reads, 650,593,822 were retained for analysis. Custom perl scripts were then used to remove barcode sequences from reads and to match reads to the beetle from which they came using their respective barcodes. Once reads were assigned to their source beetle, they were aligned to the *C. maculatus* reference genome again using the bwa aln algorithm with the same settings used for PhiX alignment. Across samples, 99,336,635 reads mapped to the reference genome constituting 65% (s.d. = 4.96%) of reads per beetle. Variant calling was performed to identify sites with single nucleotide polymorphisms (SNPs) using the bcftools call function (version 1.16) [83]. This was done using the original consensus caller (-c), retaining only variants for which the posterior probability of being invariant was less than 0.01 (-p = 0.01). Custom perl scripts were used to filter variable sites to exclude sites with a phred-scale mapping quality lower than 30, coverage less than 768 (2× the number of samples), fewer than 10 reads for the alternate allele, and with more than 20% missingness in samples (i.e., more than 20% of samples with 0 reads for a SNP). SNPs with base-quality rank-sum, mapping-quality rank-sum, or read-position rank-sum test *P* −values less than 0.001, 0.0001, and 0.001 were not retained. An additional set of filtering was performed in which SNPs with a read depth exceeding 5,030 (3 standard deviations from the mean) were removed to remove potential paralogs and gene families. Variable sites less than 2 base pairs apart were also removed. Quality filtering resulted in 11,946 SNPs for downstream analysis.

Genotype estimates were inferred from genotype likelihoods using entropy (version 2.0) [92, 93]. These Bayesian estimates integrate uncertainty in genotype likelihoods with prior information about ancestry to produce posterior genotype probabilities. The use of population models incorporating ancestry information has been shown to improve estimates of genotypes from sequence data relative to methods that do not, especially at sites with low coverage [94]. The Markov-Chain Monte Carlo (MCMC) simulation was run with 10,000 steps and a 5,000 step burn-in. The model ran under the expectation of three putative populations based on three different geographic origins of seed beetle chosen for the BILs. Allele frequencies (*p*) were calculated from genotype estimates (*g_ij_*) across *n* individuals at each SNP (*j*) using:

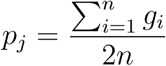

To identify regions with site-frequency spectra potentially consistent with non-netural processes during long-term maintenance on cowpea, Tajima’s *D* was calculated from the filtered vcf file. This was performed in 10 kb sliding windows with a 1 kb step size using the TajimaD function in vcftools (version 0.1.15) for the cowpea-adapted ancestors of each BIL [95]. *D* was summarized in 5 Mb rolling windows for visual comparison to other patterns of genetic differentiation (Figure S5). Then, allele frequencies were calculated using genotype estimates, from which Nei’s *F_st_*[96] was estimated between cowpea ancestors and their descendant BILs. *F_ST_* was estimated in 10 kb non-overlapping windows using the formula:

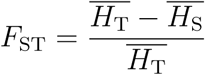

in which *H*_T_ represents total expected heterozygosity from the combined allele frequency of each population pair, and *H*_S_ represents within-population heterozygosity.

To identify regions of the genome that were most resistant to introgression in the BILs, we estimated effective migration rates (*m_e_*), or the actual rate of genetic influx into a population following migration, using an Approximate Bayesian Computation (ABC) approach [97]. Because the expected migration rate was determined by controlled crosses, deviation from this rate suggests other processes affected evolution, such as selection. For each SNP, evolution was simulated 1 million times using a forward-in-time deterministic model following the formula:

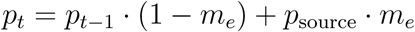

in which prior values of *m_e_* were selected from a uniform distribution from 0 to 0.5 [98]. This range was chosen as 0.5 is the migration rate established through controlled crosses during backcrossing, and therefore likely represents an upper bounds of the effective migration rate (*m_e_*). This was applied for each generation in which introgression was experienced (generations 1, 2, and 5, respectively). For all seven generations, a stochastic component approximated by binomial sampling given contemporary effective population size (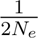, estimated using varne as done previously) was integrated into the model to account for genetic drift. SNPs whose allele frequencies differed by more than 0.1 between cowpea and lentil-adapted lines were retained for ABC simulations. These sites were presumably more informative for ancestry inference because larger allele frequency differences between populations provided greater power to detect allele frequency changes attributable to migration. The number of retained SNPs averaged 1,812.26 (s.d. = 615.06), ranging from 1,290 to 2,647 across lines. Posterior distributions for *m* were summarized using the package abc in R (version 2.2.2) [99], which retained the 100 simulations of the original 1 million (following a tolerance of 0.01%) in which the simulated final allele frequency most closely matched observed final allele frequency values.

### Early-stage adaptation experiment

To identify genetic variation that was segregating during the early stages of adaptation, two replicate lentil lines were established from a single source BF line in the same manner as the previous adaptation experiment. DNA was extracted from female beetles of the founder cowpea line as well as from the lentil-adapted lines. Both lines (BF3 and BF4) were sampled during the first generation, and BF4 was additionally sampled in the second generation to maximize the chance of capturing adaptive variation segregating at intermediate frequencies. The protocol for sequencing and processing of beetles for GBS is the same as was followed for the BILs. Ninety-six beetles from each line × generation treatment were sequenced from each line, using an Illumina Nextseq 2000 at the Genomics Core at Utah State University. Sequencing returned 842,780,472 150 bp reads, and PhiX filtering left 686,018,875 reads for downstream analysis. After trimming polyG tails, alignment was conducted following the same alignment protocol used for the BILs, producing 203,571,153 successfully aligned reads (25.1%, s.d. = 10.0% per beetle) at approximately 20× coverage. This high standard deviation is predominantly explained by a bimodal distribution of alignment success. The mean and s.d. for the two lines with poorer mapping (BF1 and BF3 F_1_) was 17.6% (s.d. = 5.5%), and 32.7% (s.d. = 7.6%) for the two lines with better mapping (BF4 F_1_ and BF4 F_2_). Still, 17,405 high-quality SNPs were obtained based on the same strict variant filtering guidelines used in the BILs. Genotypes were then estimated using the same entropy settings as above, but with 20,000 steps and a 10,000-step burn-in [93].

Regions of local population structure underlying putative structural variation were identified using a windowed principal component analysis (PCA) approach [47]. For this, WinPCA (version 1.2) used genotype estimates to generate local PCAs for 10 Mb windows in 10 kb sliding steps [100]. This window size was chosen to be the same order of magnitude as the regions of parallel differentiation (RPDs) in the whole genome dataset. Sliding window principal component analyses were run separately for each lentil line alongside the founder line to prevent population-specific structure from confounding signatures of local structure created by reduced recombination. BF3 F_1_ was used for Figure 3 as it was the best visual representative (but see Figure S1). Hierarchical clustering was performed in each PC window using the hclust and cutree functions from the stats package in R (version 4.4.0) to identify clusters in the first principal component (*k* = 3). Regions exhibiting structure were identified using the Calinski-Harabasz (CH) index [101], implemented using the calinhara function from the fpc package (version 2.2-14) in R [102]. Given *n* points and *k* clusters, this index evaluates clustering quality as the ratio of the between-cluster separation (*BCSS*) to the within-cluster dispersion (*WCSS*), controlled by their degrees of freedom such that:

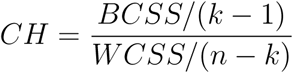

A clustering coefficient of *k* = 3 was chosen as this mirrors expectations under three segregating Mendelian genotypes, as would be the case with simple structural variants (i.e., inversions). Regions of putative structure on chromosomes 1 and 9 were chosen based on elevated CH index values within the RPDs which were identified as involved in adaptation in the BILs. Within these two regions (chr1:77-86 Mb and chr9:47-62 Mb respectively), individual inbreeding coefficients (F_IS_) were calculated using vcftools (version 0.1.15) for each individual along the bounds of large, segregating haploblocks. Kruskal-Wallis tests were performed on F_IS_ between each cluster using the kruskal.test function in R to assess whether significant differences between clusters for inbreeding values served as a proxy for genotype delineation. Finally, pairwise linkage disequilibrium (LD) was estimated using PLINK (version v1.90b5.3) for chromosomes 1 and 9 [103], which was then aggregated into 1 Mb windows for visualization in R.

### Differential Expression Analysis

In Rego et al. [45], three separate adaptation lines of *C. maculatus* of the South India (SI) lineage were established from mung (an equally suitable host to cowpea) lines shifted to lentil to both estimate patterns of differential expression between host treatments and to map the genetic basis of fitness components (see ‘Genome-wide Association Mapping’). The experimental design and culture procedures for this study follow those for previous host-shift experiments. Briefly, RNA was extracted from replicate lines created from SI-L1, which had been maintained on lentil for 107 generations at the time of sampling, and from SI-M, which had been maintained on mung for >300 generations. Samples were taken from each line on their current host, as well as each line following a generation on the alternate host when survival was possible (mung-adapted beetles did not produce enough offspring on lentil for sufficient sampling). Fourth instar (final instar) larvae were used as this developmental period is characterized by active feeding, so exposure to the effects of a novel nutritional regime was likely at its highest.

Larvae were flash-frozen and prepared for extraction in the pre- and post-adapted lines. Total RNA from five biological replicates of three pooled 4th-instar larvae for each treatment was isolated using RNeasy Mini kits (Qiagen Inc.). After quality control and library preparation, RNAseq libraries were quantitated using qPCR and sequenced on a HiSeq2500 (Illumina, USA). This project generated 880,655,484 reads with a mean and s.d. of 35,226,219 (s.d. = 3,061,575) per library, and the sequence data is available on NCBI’s SRA database (accession PRJNA616195). Raw reads were processed in Rego et al. [45], for which RCorrector (version v1.0.4) was used to fix k-mer based read errors [104], and a custom python script was used to remove uncorrected reads. Reads were then trimmed to remove Illumina adapter sequences using TrimGalore! (version 0.3.3) [105]. In the current study, these trimmed, high quality paired-end reads were aligned to the updated chromosome-level reference genome using STAR (version 2.7.10a) [106]. The percent mean and s.d. of reads which aligned to the reference genome across samples was 72.9%, s.d. = 2.5% Finally, featureCounts (version 2.0.0) was used to convert STAR alignments to gene count data [107].

Filtering, normalization, and differential expression analyses followed the same protocol as Rego et al. [45]. Here, we filtered and normalized count data using the edgeR library in R (version 4.10.7) [108]. Genes with low expression were removed using edgeR, after normalizing count data through estimation of counts per million (CPM) values for each gene using the cpm function. Specifically, genes with CPM values > 0.5 in at least two samples were retained for differential expression analyses, leaving 8,237 genes. Differential expression analyses were carried out using the limma library in R (version 3.68.5) [109]. First, variance stabilizing normalization was conducted with voom [110], using a design matrix corresponding to the linear model used for the analysis. The model.matrix function was used to fit model design matrices using the host and line as factor combinations, and the makeContrasts function was used to extract comparisons of interests as contrasts. The significance of model effects was calculated using voom precision weights and retained effects. The decideTests function was used to determine if a model effect was significant and retained effects if their Benjamini-Hochberg false-discovery rate corrected *P*-value was less than 0.05. For this work, contrasts were only made between mung-adapted beetles reared on mung, and lentil-adapted beetles reared on mung. This was to ensure that differences in expression were the result of genetic differences rather than environmental differences (i.e., plasticity). Mung was chosen as the common host due to the lack of surviving mung-adapted beetles on green lentil. The contrasts estimated with voom were used to identify genes which showed significant difference in gene expression. Finally, permutation tests were run to identify RPDs which had more differentially expressed genes within the 99th percentile of − log_10_ *P*-values than expected based on genomic windows of similar size.

### Genome-wide Association Mapping

In addition to differential expression analyses, Rego et al. [45] created backcross lines from SI lentil-adapted lines to map the genetic basis of body mass and development time, two highlevel fitness components. Backcross mapping was conducted on three SI lines (SI-L1, SI-L2, and SI-L14A), which had been adapted to lentil for 146, 135, and 38 generations respectively. To accomplish this, F_1_ hybrids were generated by crossing each lentil-adapted line with an SI line maintained on their ancestral host (i.e., mung bean). The resulting F_1_ individuals were then backcrossed to the corresponding lentil-adapted line. Across all three mapping lines, time to adult emergence and adult body mass were measured in 748 female beetles (241, 256, and 251 from L1, L2, and L14A respectively). These beetles were sequenced using the same GBS protocol described above, and the same bioinformatics pipeline was used to produce high quality genotype estimates for trait mapping and other downstream analyses, with alignment percentages of 61%, s.d. = 4.18% (mean and s.d.) across samples. Here, the Markov-Chain Monte Carlo (MCMC) for entropy was run with 20,000 steps and a 10,000-step burn-in to produce genotype estimates [93]. This workflow resulted in 27,931 high quality SNPs for trait mapping and other downstream analyses.

These SNPs were used to infer the genetic contribution to adult mass and development time in the backcrossed lines. We accomplished this using a Bayesian sparse linear mixed model (BSLMM) implemented in gemma (version 0.98.3) [111]. This method differs from traditional genome-wide association (GWA) methods in that it models the effect of all SNPs simultaneously, generating an estimate of the polygenic contribution of each SNP to trait associations (*α*) to avoid the pitfall of multiple hypothesis testing [111]. The polygenic effect determines the collective contribution of all SNPs on phenotypic variance, and accounts for phenotypic covariances among individuals as a result of genetic similarity (i.e., observed kinship). For each SNP, gemma estimates a polygenic effect (*α*), a posterior inclusion probability (PIP; *γ*) for the sparse component, and its effect conditional on inclusion (*β*); the resulting total effect is *α* + *βγ* [112]. The total effect size of a given SNP can then be determined by the sum of the polygenic and sparse parameters conditional on the posterior inclusion probability (*α_i_*+ *β_i_γ_i_*).

This hierarchical structure allows for the estimation of additional parameters that are useful for understanding a trait’s genetic architecture, such as the percent phenotypic variance explained by additive effects (PVE) and the percent of genetic variance explained by SNPs with measurable effects (PGE, *β* ≠ 0). The sparse component is expected to count only a single causal variant when multiple SNPs share similar genotype trajectories, making its estimates generally robust to LD. Here, the BSLMM was run separately for each line × phenotype combination with ten chains and the minor allele frequency filter (-maf) set to 0.001. The Markov-Chain Monte Carlo (MCMC) was run with a burn-in (-w) of 1 million steps, after which the model ran with an additional 1 million steps (-s). The state of every 20th step was recorded (-rpace), leaving 50,000 observations from which posteriors were constructed.

To further understand the genetic basis of adaptive trait variation, local genetic structure was visualized using WinPCA [100] in similar fashion to the early-sequenced experiment with 10 Mb overlapping windows of local PC structure estimated in 10 kb steps. PC1 scores for individual beetles were colored corresponding to trait value for both development time and adult mass. While all lines were visualized in this manner (Figures S2-S4), the first PC of adult mass in L1 was included in Figure 5 due to the most visually clear segregation of genotypes in the 71-85 Mb region. A hierarchical clustering analysis at 79 Mb was used to determine which individuals clustered into the putative haplotype group in each line as this region visually exhibited strongest structure. vcftools (version 0.1.15) was then used to again estimate individual inbreeding coefficients (*F*) between 71-85 Mb, and Kruskal-Wallis tests were conducted using the kruskal.test function in R to determine whether the clusters differed significantly for inbreeding, body mass, and development time respectively. Finally, pairwise linkage disequilibrium (LD) was again estimated using PLINK and aggregated into 1 Mb windows in R [103].

## Supplemental Figures and Tables

**Table S1:**
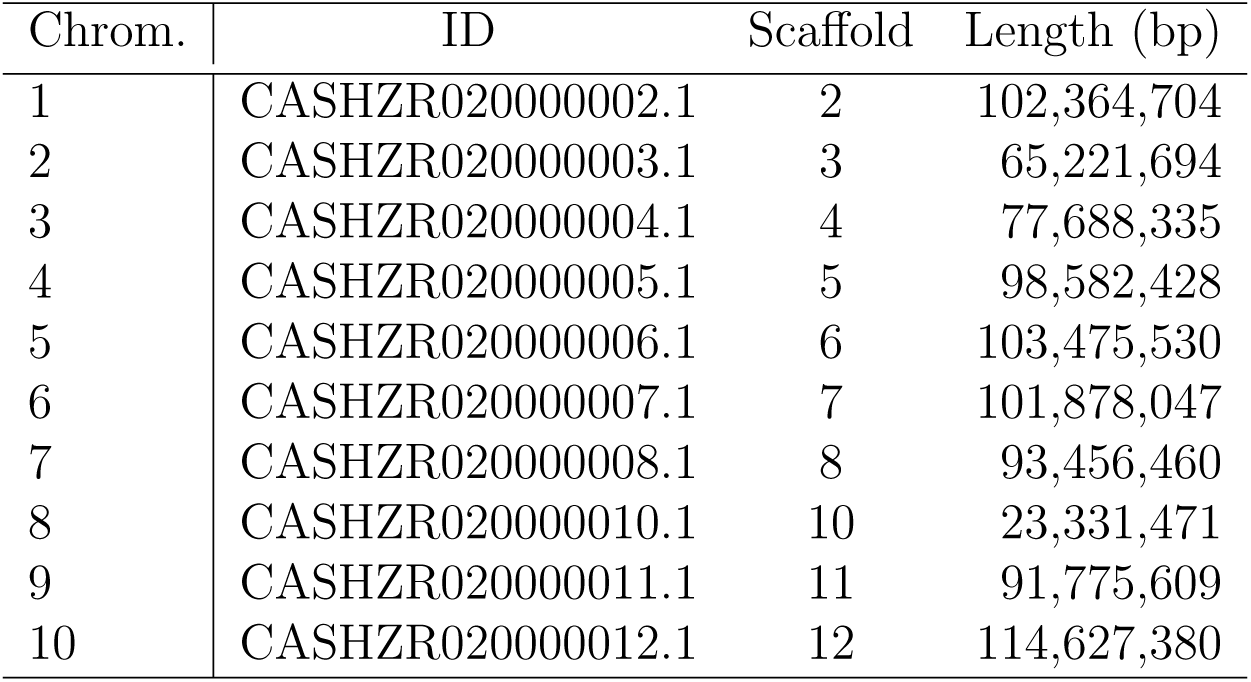
Chromosome numbering system used in this paper and corresponding scaffold numbers from [46]. Chrom. provides our chromosome numbers, ID denotes the genome fasta headers, scaffold reports the scaffold numbers used in [46] (see their Figure 2 in particular, and length reports the scaffold lengths in base pairs (bp). Our chromosome 8 is likely the X sex chromosome.

| Chrom. | ID | Scaffold | Length (bp) |
|---|---|---|---|
| 1 | CASHZR020000002.1 | 2 | 102,364,704 |
| 2 | CASHZR020000003.1 | 3 | 65,221,694 |
| 3 | CASHZR020000004.1 | 4 | 77,688,335 |
| 4 | CASHZR020000005.1 | 5 | 98,582,428 |
| 5 | CASHZR020000006.1 | 6 | 103,475,530 |
| 6 | CASHZR020000007.1 | 7 | 101,878,047 |
| 7 | CASHZR020000008.1 | 8 | 93,456,460 |
| 8 | CASHZR020000010.1 | 10 | 23,331,471 |
| 9 | CASHZR020000011.1 | 11 | 91,775,609 |
| 10 | CASHZR020000012.1 | 12 | 114,627,380 |

**Table S2:** Differentially transcribed genes within RPD:4 (chr1:54-95 Mb) exhibit variable putative functions.

| **Gene** | **logFC** | **FDR *P*-value** | **Accession** | **Name** | **Function** |
|---|---|---|---|---|---|
| g4556 | 1.7908 | 3.08E-03 | XM_023173976.2 | Putative divalent cation/proton antiporter TMEM165 | Golgi calcium homeostasis |
| g4573 | 0.6716 | 6.03E-03 | XM_020023651.2 | Eukaryotic translation initiation factor 5B-like | GTP binding, translation initiation |
| g4600 | 0.8429 | 7.15E-03 | XM_018714107.2 | Tectonin beta-propeller repeat-containing protein | Carbohydrate binding, membrane |
| g4615 | -3.1361 | 7.96E-03 | XM_018715671.1 | Alpha-tocopherol transfer protein-like | Phosphatidylinositol bisphosphate binding |
| g4634 | 4.7415 | 1.54E-03 | XM_023162095.2 | Splicing factor C9orf78 | mRNA splicing via spliceosome |
| g4639 | 1.4799 | 4.01E-04 | XM_088035195.1 | Grindelwald (grnd) | TNF receptor activity, cell death / ROS reg |
| g4645 | 0.9523 | 1.30E-03 | - | - | - |
| g4659 | 0.6193 | 5.55E-03 | XM_056787621.1 | Protein Malvolio-like | Cadmium transport, manganese homeostasis |
| g4711 | -5.4545 | 2.63E-05 | XR_007664227.1 | Non-structural maintenance of chromosomes 4A | Smc5-Smc6 complex, DNA repair |
| g4763 | 3.6749 | 6.90E-03 | XM_023156345.2 | Uncharacterized protein (LOC111502292) | - |
| g4768 | -1.8941 | 6.03E-03 | XM_018709359.1 | Glyoxylate reductase/hydroxypyruvate reductase | Hydroxypyruvate reductase activity |
| g4769 | -4.6901 | 4.01E-04 | XM_023163689.2 | UbiA prenyltransferase domain-containing heix | Prenyltransferase, Vitamin K biosynthesis |
| g4855 | 0.5095 | 8.50E-03 | XM_081979322.1 | Venom serine carboxypeptidase | Serine-type carboxypeptidase, proteolysis |

**Figure S1:**
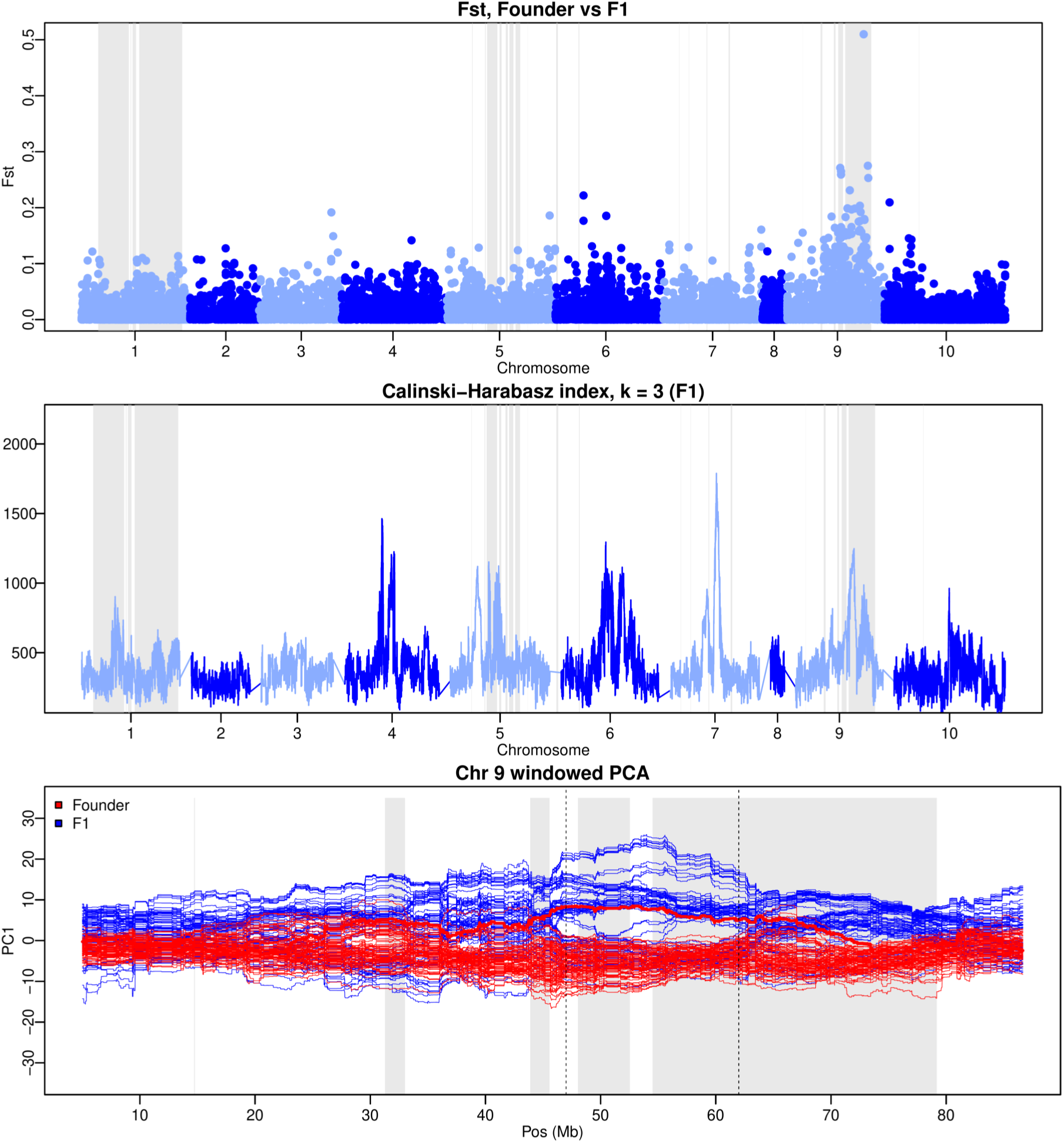
A large haploblock underlies BF4, similarly to BF3. Genome-wide F_ST_ and Calinksi-Harabasz index (*k* = 3) for BF4 F_1_, along with local PCA on Chromosome 9. Black dotted lines denote the same region used for cluster analysis in BF3 F_1_.

**Figure S2:**
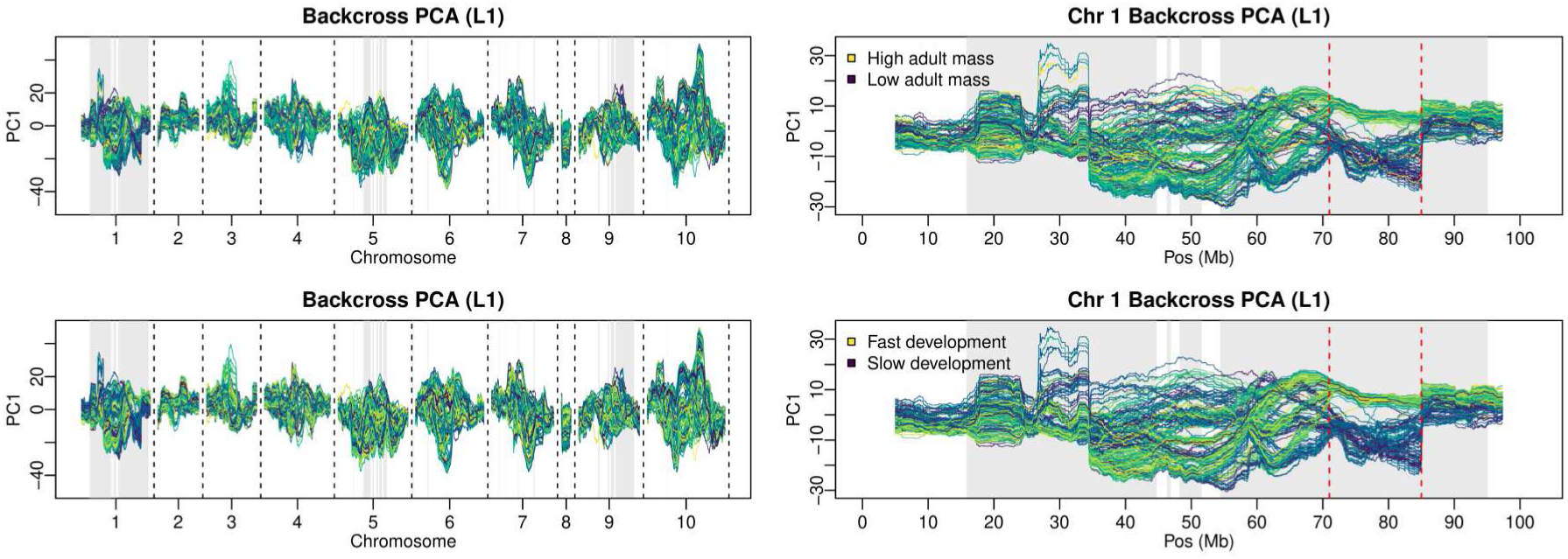
Local PCA for L1, colored by fitness component traits. Genome-wide (left) and Chromosome-1 (right) local PCA for the L1 backcross population. Individuals are colored based on trait values for adult mass (top) and development time (bottom).

**Figure S3:**
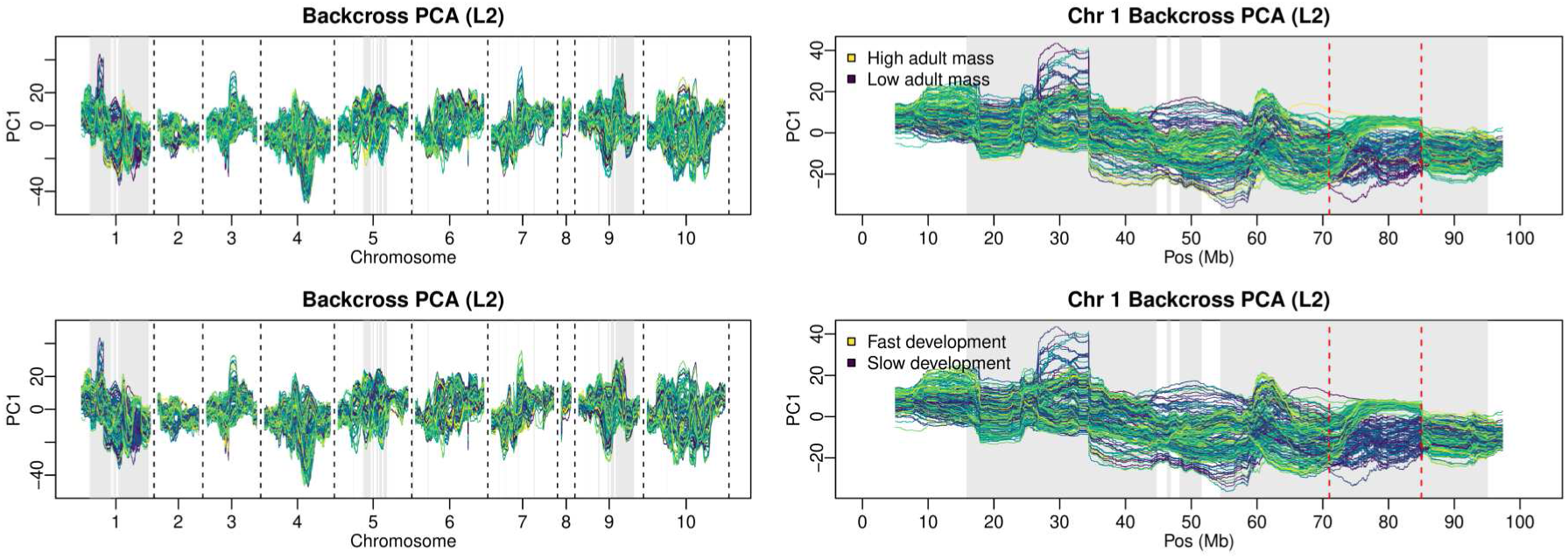
Local PCA for L2, colored by fitness component traits. Genome-wide (left) and Chromosome-1 (right) local PCA for the L2 backcross population. Individuals are colored based on trait values for adult mass (top) and development time (bottom).

**Figure S4:**
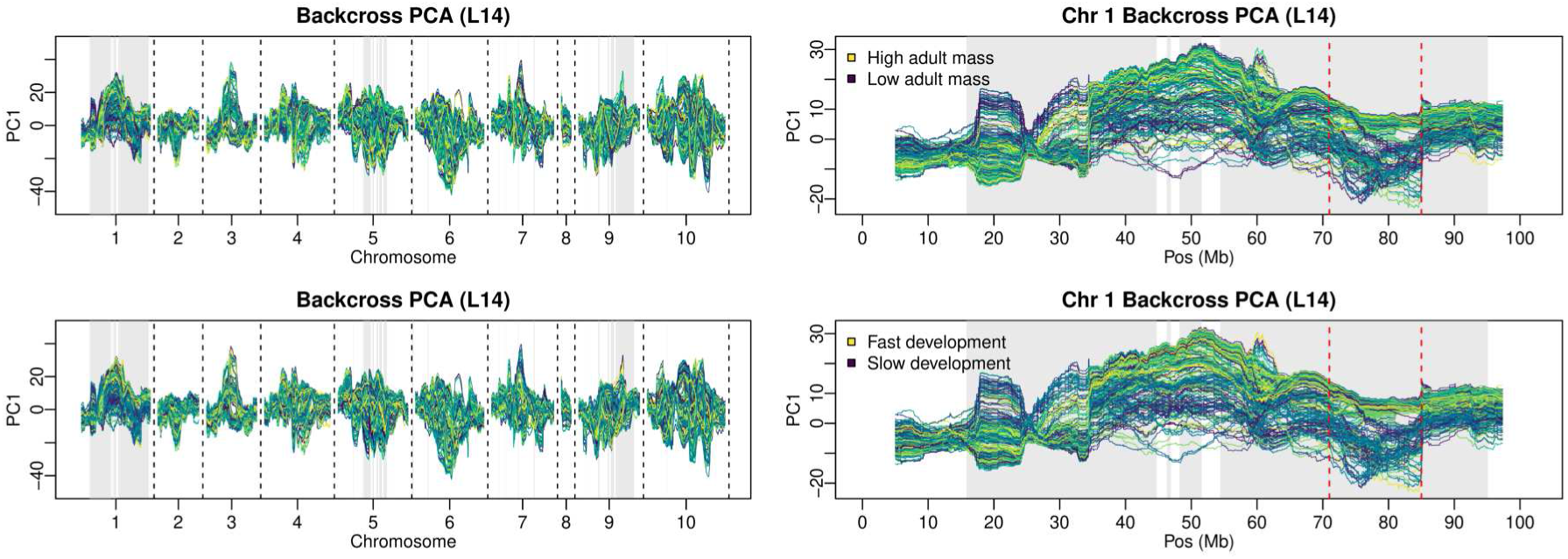
Local PCA for L14, colored by fitness component traits. Genome-wide (left) and Chromosome-1 (right) local PCA for the L14 backcross population. Individuals are colored based on trait values for adult mass (top) and development time (bottom).

**Figure S5:**
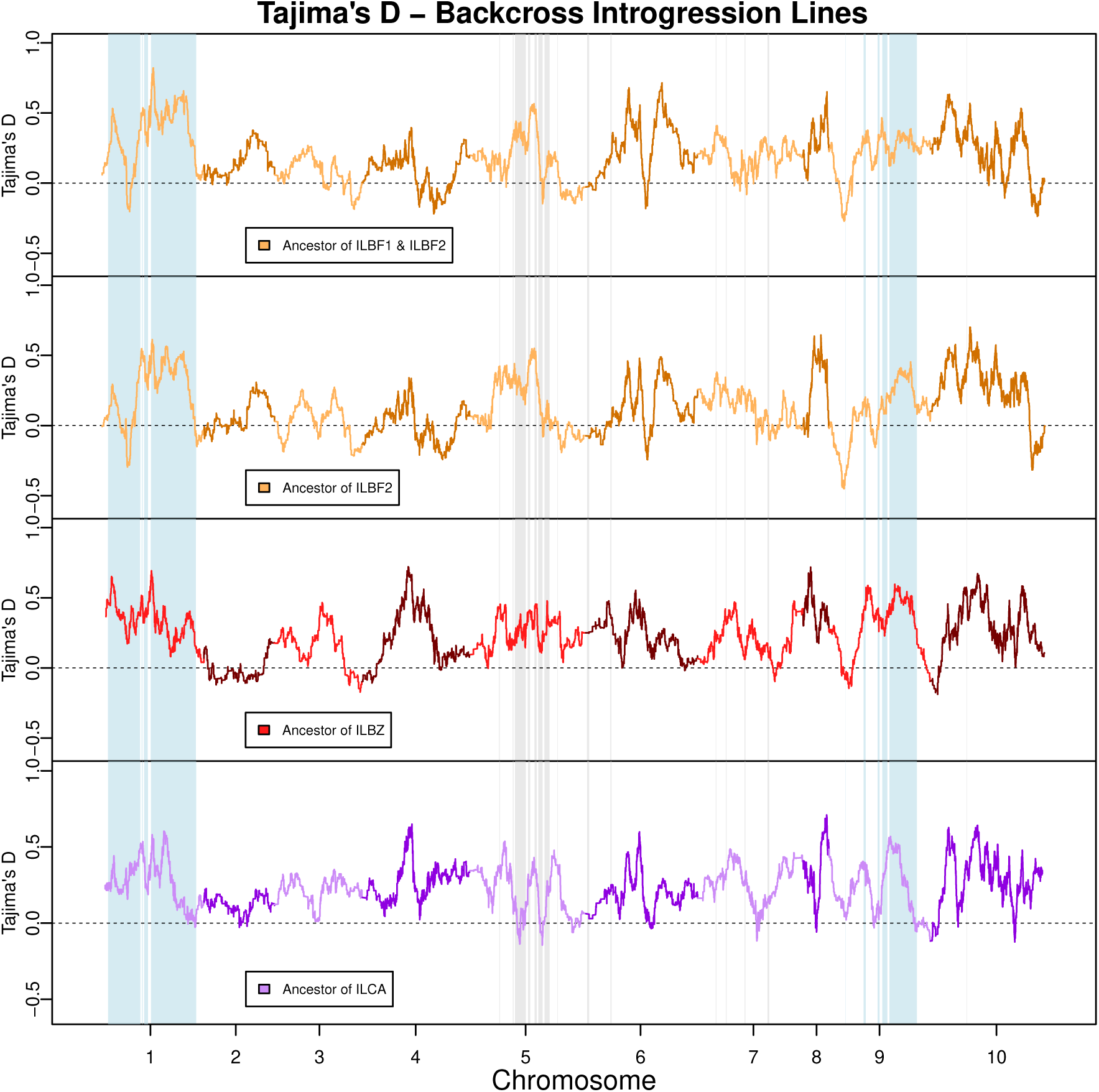
Tajima’s *D* in the cowpea-adapted ancestors of the backcross introgression lines. *D* was estimated in 10 kb non-overlapping windows, which were summarized into 5 Mb rolling windows for visualization.

## Literature Cited

[1] Barghi, N., Hermisson, J. & Schlotterer, C. Polygenic adaptation: a unifying frame-work to understand positive selection. Nature Reviews Genetics 21, 769–781 (2020).

[2] Kardos, M. & Luikart, G. The genetic architecture of fitness drives population viability during rapid environmental change. The American Naturalist 197, 511–525 (2021).

[3] Whiting, J. R. et al. On the genetic architecture of rapidly adapting and convergent life history traits in guppies. Heredity 128, 250–260 (2022).

[4] Bomblies, K. & Peichel, C. L. Genetics of adaptation. Proceedings of the National Academy of Sciences 119, e2122152119 (2022).

[5] Souaiaia, T. et al. Distinct genetic architecture in the tails of complex traits. Nature 1–9 (2026).

[6] Tigano, A. & Friesen, V. L. Genomics of local adaptation with gene flow. Molecular Ecology 25, 2144–2164 (2016).

[7] Taylor, C. S. & Lawson, D. J. Heritability of complex traits in sub-populations experiencing bottlenecks and growth. Journal of Human Genetics 69, 329–335 (2024).

[8] O’Connor, L. J. The distribution of common-variant effect sizes. Nature Genetics 53, 1243–1249 (2021).

[9] Yeaman, S. Evolution of polygenic traits under global vs local adaptation. Genetics 220, iyab134 (2022).

[10] Kreiner, J. M. The genetic architecture and spatiotemporal dynamics of adaptation across human-modified landscapes. New Phytologist 249, 744–750 (2026).

[11] Orr, H. A. & Unckless, R. L. The population genetics of evolutionary rescue. PLOS Genetics 10, e1004551 (2014).

[12] Gomulkiewicz, R. & Holt, R. D. When does evolution by natural selection prevent extinction? Evolution 201–207 (1995).

[13] Balstad, L. J., Baskett, M. L., Carlson, S. M., Cunningham, C. J. & Westley, P. A. Revisiting evolutionary rescue in the wild. Trends in Ecology & Evolution.

[14] Heinen, R., Thakur, M. P., Harvey, J. A. & Gols, R. Effects of extreme climatic events on evolutionary processes. Global Change Biology 31, e70618 (2025).

[15] Sparks, M., Schraidt, C., Yin, X., Seeb, L. & Christie, M. Rapid genetic adaptation to a novel ecosystem despite a massive bottleneck (2023).

[16] Henschen, A. E. et al. Rapid adaptation to a novel pathogen through disease tolerance in a wild songbird. PLoS Pathogens 19, e1011408 (2023).

[17] Dogantzis, K. A. et al. Post-invasion selection acts on standing genetic variation despite a severe founding bottleneck. Current Biology 34, 1349–1356 (2024).

[18] Charlesworth, B. Effective population size and patterns of molecular evolution and variation. Nature Reviews Genetics 10, 195–205 (2009).

[19] Gravel, S. When is selection effective? Genetics 203, 451–462 (2016).

[20] Kimura, M. On the probability of fixation of mutant genes in a population. Genetics 47, 713 (1962).

[21] Dlugosch, K. M., Anderson, S. R., Braasch, J., Cang, F. A. & Gillette, H. D. The devil is in the details: genetic variation in introduced populations and its contributions to invasion. Molecular Ecology 24, 2095–2111 (2015).

[22] Yeaman, S. & Whitlock, M. C. The genetic architecture of adaptation under migration– selection balance. Evolution 65, 1897–1911 (2011).

[23] Kirkpatrick, M. & Barton, N. Chromosome inversions, local adaptation and speciation. Genetics 173, 419–434 (2006).

[24] Villoutreix, R. et al. Inversion breakpoints and the evolution of supergenes. Molecular Ecology 30, 2738–2755 (2021).

[25] Olazcuaga, L., Melbourne, B. A., Nordstrom, S. W. & Hufbauer, R. A. Density dependence during evolutionary rescue increases extinction risk but does not prevent adaptation. Ecology Letters 29, e70312 (2026).

[26] Leigh, D. M., Lischer, H. E., Guillaume, F., Grossen, C. & Gunther, T. Disentangling adaptation from drift in bottlenecked and reintroduced populations of alpine ibex. Molecular Ecology Resources 21, 2350–2363 (2021).

[27] Kruglyak, L. Prospects for whole-genome linkage disequilibrium mapping of common disease genes. Nature Genetics 22, 139–144 (1999).

[28] Xenikoudakis, G. et al. Consequences of a demographic bottleneck on genetic structure and variation in the scandinavian brown bear. Molecular Ecology 24, 3441–3454 (2015).

[29] Gompert, Z., Feder, J. L. & Nosil, P. Natural selection drives genome-wide evolution via chance genetic associations. Molecular Ecology 31, 467–481 (2022).

[30] Bailey, S. F. & Bataillon, T. Can the experimental evolution programme help us elucidate the genetic basis of adaptation in nature? Molecular Ecology 25, 203–218 (2016).

[31] Kawecki, T. J. et al. Experimental evolution. Trends in Ecology & Evolution 27, 547–560 (2012).

[32] Santos, M. A. et al. Experimental evolution in a warming world: the omics era. Molecular Biology and Evolution 41, msae148 (2024).

[33] Kassen, R. Toward a functional genetics of adaptation: Insights from microbial experimental evolution. Genome Biology and Evolution 18, evag158 (2026).

[34] Graves Jr, J., et al. Genomics of parallel experimental evolution in *Drosophila*. Molecular Biology and Evolution 34, 831–842 (2017).

[35] Baldwin-Brown, J. G., Long, A. D. & Thornton, K. R. The power to detect quantitative trait loci using resequenced, experimentally evolved populations of diploid, sexual organisms. Molecular Biology and Evolution 31, 1040–1055 (2014).

[36] Long, A., Liti, G., Luptak, A. & Tenaillon, O. Elucidating the molecular architecture of adaptation via evolve and resequence experiments. Nature Reviews Genetics 16, 567–582 (2015).

[37] Nagata, K. et al. Series of advanced backcross populations uncovers diversity and complexity in genetic control of grain length and grain width in asian cultivated rice. Breeding Science 76, 251–262 (2026).

[38] Zhang, B., Ma, L., Wu, B., Xing, Y. & Qiu, X. Introgression lines: Valuable resources for functional genomics research and breeding in rice (*Oryza sativa L.*). Frontiers in Plant Science 13, 863789 (2022).

[39] Messina, F. J., Lish, A. M. & Gompert, Z. Variable responses to novel hosts by populations of the seed beetle *Callosobruchus maculatus* (Coleoptera: Chrysomelidae: Bruchinae). Environmental Entomology 47, 1194–1202 (2018).

[40] Rego, A., Messina, F. J. & Gompert, Z. Dynamics of genomic change during evolutionary rescue in the seed beetle *Callosobruchus maculatus*. Molecular Ecology 28, 2136–2154 (2019).

[41] Springer, A., Kissmer, B. & Gompert, Z. Admixture affects the rate and repeatability of experimental adaptation to a stressful environment in *Callosobruchus maculatus*. Molecular Ecology 34, e17809 (2025).

[42] Kebe, K., et al. Global phylogeography of the insect pest *Callosobruchus maculatus* (Coleoptera: Bruchinae) relates to the history of its main host, *Vigna unguiculata*. Journal of Biogeography 44, 2515–2526 (2017).

[43] Messina, F. J., Lish, A. M., Springer, A. & Gompert, Z. Colonization of marginal host plants by seed beetles (Coleoptera: Chrysomelidae): Effects of geographic source and genetic admixture. Environmental Entomology 49, 938–946 (2020).

[44] Messina, F. J., Jones, J. C., Mendenhall, M. & Muller, A. Genetic modification of host acceptance by a seed beetle, *Callosobruchus maculatus* (Coleoptera: Bruchidae). Annals of the Entomological Society of America 102, 181–188 (2009).

[45] Rego, A., et al. Combining experimental evolution and genomics to understand how seed beetles adapt to a marginal host plant. Genes 11, 400 (2020).

[46] Arnqvist, G., et al. A chromosome-level assembly of the seed beetle *Callosobruchus maculatus* genome with annotation of its repetitive elements. G3 Genes|Genomes|Genetics 14, jkad266 (2023).

[47] Li, H. & Ralph, P. Local pca shows how the effect of population structure differs along the genome. Genetics 211, 289–304 (2019).

[48] Buffalo, V. & Coop, G. Estimating the genome-wide contribution of selection to temporal allele frequency change. Proceedings of the National Academy of Sciences 117, 20672–20680 (2020).

[49] Kissmer, B. & Gompert, Z. Chromosomal rearrangements might play a central role in adaptation. Molecular Ecology 34, e70159 (2025).

[50] Gompert, Z. et al. Adaptation repeatedly uses complex structural genomic variation. Science 388, eadp3745 (2025).

[51] Huang, K. et al. Inversions contribute disproportionately to parallel genomic divergence in dune sunflowers. Nature Ecology & Evolution 9, 325–335 (2025).

[52] Villoutreix, R. et al. Large-scale mutation in the evolution of a gene complex for cryptic coloration. Science 369, 460–466 (2020).

[53] Thurman, T. J. & Barrett, R. D. The genetic consequences of selection in natural populations. Molecular Ecology 25, 1429–1448 (2016).

[54] Messina, F. J. & Durham, S. L. Loss of adaptation following reversion suggests tradeoffs in host use by a seed beetle. Journal of Evolutionary Biology 28, 1882–1891 (2015).

[55] McAllester, C. S. & Pool, J. E. The potential of inversions to accumulate balanced sexual antagonism is supported by simulations and drosophila experiments. Elife 12, RP93338 (2025).

[56] Paris, M. et al. Multiple forms of balancing selection maintain inversion polymorphism. Heredity 135, 138–151 (2026).

[57] Ruzicka, F. et al. A century of theories of balancing selection. Biological Reviews 101, 804–825 (2026).

[58] Dougherty, L. R. et al. Sexual conflict and correlated evolution between male persistence and female resistance traits in the seed beetle *Callosobruchus maculatus*. Proceedings of the Royal Society B: Biological Sciences 284, 20170132 (2017).

[59] Edvardsson, M. & Canal, D. The effects of copulation duration in the bruchid beetle *Callosobruchus maculatus*. Behavioral Ecology 17, 430–434 (2006).

[60] Eady, P. E., Hamilton, L. & Lyons, R. E. Copulation, genital damage and early death in *Callosobruchus maculatus*. Proceedings of the Royal Society B: Biological Sciences 274, 247–252 (2007).

[61] Berdan, E. L. et al. How chromosomal inversions reorient the evolutionary process. Journal of Evolutionary Biology 36, 1761–1782 (2023).

[62] Faria, R., Johannesson, K., Butlin, R. K. & Westram, A. M. Evolving inversions. Trends in Ecology & Evolution 34, 239–248 (2019).

[63] Messina, F. J., Mendenhall, M. & Jones, J. C. An experimentally induced host shift in a seed beetle. Entomologia Experimentalis et Applicata 132, 39–49 (2009).

[64] Chaturvedi, S. et al. Climatic similarity and genomic background shape the extent of parallel adaptation in *Timema* stick insects. Nature Ecology & Evolution 6, 1952–1964 (2022).

[65] Bohutınska, M. & Peichel, C. L. Divergence time shapes gene reuse during repeated adaptation. Trends in Ecology & Evolution 39, 396–407 (2024).

[66] Owens, G. L. et al. Shared selection and genetic architecture drive strikingly repeatable evolution in long-term experimental hybrid populations. Molecular Biology and Evolution msaf014 (2025).

[67] Schlotterer, C. How predictable is adaptation from standing genetic variation? experimental evolution in *Drosophila* highlights the central role of redundancy and linkage disequilibrium. Philosophical Transactions of the Royal Society B: Biological Sciences 378, 20220046 (2023).

[68] Johnson, M. S., Reddy, G. & Desai, M. M. Epistasis and evolution: recent advances and an outlook for prediction. BMC Biology 21, 120 (2023).

[69] Tuda, M., Kagoshima, K., Toquenaga, Y. & Arnqvist, G. Global genetic differentiation in a cosmopolitan pest of stored beans: effects of geography, host-plant usage and anthropogenic factors. PLoS One 9, e106268 (2014).

[70] Tuda, M., Ronn, J., Buranapanichpan, S., Wasano, N. & Arnqvist, G. Evolutionary diversification of the bean beetle genus *Callosobruchus* (Coleoptera: Bruchidae): traits associated with stored-product pest status. Molecular Ecology 15, 3541–3551 (2006).

[71] Larson, A. O. & Fisher, C. K. The bean weevil and the southern cowpea weevil in California (1938).

[72] Credland, P. F., Dick, K. M. & Wright, A. W. Relationships between larval density, adult size and egg production in the cowpea seed beetle, Callosobruchus maculatus. (1986).

[73] Messina, F. J. Heritability and ‘evolvability’ of fitness components in *Callosobruchus maculatus*. Heredity 71, 623–629 (1993).

[74] Tran, B. M. & Credland, P. F. Consequences of inbreeding for the cowpea seed beetle, *Callosobruchus maculatus* (F.)(Coleoptera: Bruchidae). Biological Journal of the Linnean Society 56, 483–503 (1995).

[75] Fox, C. W. A quantitative genetic analysis of oviposition preference and larval performance on two hosts in the bruchid beetle, *Callosobruchus maculatus*. Evolution 47, 166–175 (1993).

[76] Dowling, D. K., Abiega, K. C. & Arnqvist, G. Temperature-specific outcomes of cytoplasmic-nuclear interactions on egg-to-adult development time in seed beetles. Evolution 61, 194–201 (2007).

[77] Bhattacharya, B., Barik, A. & Banerjee, T. Bioenergetics and water balance in *Callosobruchus maculatus* (F.)(Coleoptera: Bruchidae) larval populations. Oriental Insects 37, 423–437 (2003).

[78] Guedes, R. N. C., Smith, R. H. & Guedes, N. M. P. Host suitability, respiration rate and the outcome of larval competition in strains of the cowpea weevil, *Callosobruchus maculatus*. Physiological Entomology 28, 298–305 (2003).

[79] Li, H. & Durbin, R. Fast and accurate short read alignment with burrows–wheeler transform. Bioinformatics 25, 1754–1760 (2009).

[80] Li, H. Aligning sequence reads, clone sequences and assembly contigs with bwa-mem. arXiv preprint arXiv:1303.3997 (2013).

[81] Li, H. et al. The sequence alignment/map format and samtools. Bioinformatics 25, 2078–2079 (2009).

[82] Van der Auwera, G. A. & O’Connor, B. D. Genomics in the cloud: using Docker, GATK, and WDL in Terra (O’Reilly Media, 2020).

[83] Li, H. A statistical framework for snp calling, mutation discovery, association mapping and population genetical parameter estimation from sequencing data. Bioinformatics 27, 2987–2993 (2011).

[84] Gompert, Z. & Messina, F. J. Genomic evidence that resource-based trade-offs limit host-range expansion in a seed beetle. Evolution 70, 1249–1264 (2016).

[85] Ewens, W. J. & Ewens, W. Mathematical Population Genetics: Theoretical Introduction, vol. 27 (Springer, 2004).

[86] Gaggiotti, O. E. & Foll, M. Quantifying population structure using the F-model. Molecular Ecology Resources 10, 821–830 (2010).

[87] R Core Team. R: A Language and Environment for Statistical Computing. R Foundation for Statistical Computing, Vienna, Austria (2022). URL https://www.R-project.org.

[88] Booker, T. R., Yeaman, S. & Whitlock, M. C. Using genome scans to identify genes used repeatedly for adaptation. Evolution 77, 801–811 (2023).

[89] Benjamini, Y. & Hochberg, Y. Controlling the false discovery rate: a practical and powerful approach to multiple testing. Journal of the Royal Statistical Society: Series B (Methodological) 57, 289–300 (1995).

[90] Harte, D. S. HiddenMarkov: Hidden Markov Models. Statistics Research Associates, Wellington NZ (2025). URL https://www.statsresearch.co.nz/dsh/sslib/. R package version 1.8-14.

[91] Kaufmann, P. et al. Y-linked copy number polymorphism of target of rapamycin is associated with sexual size dimorphism in seed beetles. Molecular Biology and Evolution 40, msad167 (2023).

[92] Gompert, Z., Lucas, L. K., Fordyce, J. A., Forister, M. L. & Nice, C. C. Secondary contact between *Lycaeides idas* and *L. melissa* in the rocky mountains: extensive admixture and a patchy hybrid zone. Molecular Ecology 19, 3171–3192 (2010).

[93] Shastry, V. et al. Model-based genotype and ancestry estimation for potential hybrids with mixed-ploidy. Molecular Ecology Resources 21, 1434–1451 (2021).

[94] Clark, L. V., Lipka, A. E. & Sacks, E. J. polyrad: Genotype calling with uncertainty from sequencing data in polyploids and diploids. G3: Genes, Genomes, Genetics 9, 663–673 (2019).

[95] Danecek, P. et al. The variant call format and vcftools. Bioinformatics 27, 2156–2158 (2011).

[96] Nei, M. Analysis of gene diversity in subdivided populations. Proceedings of the National Academy of Sciences 70, 3321–3323 (1973).

[97] Beaumont, M. A. Approximate bayesian computation. Annual Review of Statistics and its Application 6, 379–403 (2019).

[98] Wright, S. Evolution in mendelian populations. Genetics 16, 97 (1931).

[99] Csillery, K., Francois, O. & Blum, M. G. B. abc: an r package for approximate Bayesian computation (abc). Methods in Ecology and Evolution (2012).

[100] Blumer, L. M., Good, J. M. & Durbin, R. Winpca: a package for windowed principal component analysis. Bioinformatics 41, btaf529 (2025).

[101] Caliński, T. & Harabasz, J. A dendrite method for cluster analysis. Communications in Statistics-theory and Methods 3, 1–27 (1974).

[102] Hennig, C. & Imports, M. Package ‘fpc’. Flexible Procedures for Clustering 1176 (2015).

[103] Purcell, S. et al. Plink: a tool set for whole-genome association and population-based linkage analyses. The American Journal of Human Genetics 81, 559–575 (2007).

[104] Song, L. & Florea, L. Rcorrector: efficient and accurate error correction for Illumina RNA-seq reads. GigaScience 4, s13742–015 (2015).

[105] Martin, M. Cutadapt removes adapter sequences from high-throughput sequencing reads. *EMBnet*. journal 17, 10–12 (2011).

[106] Dobin, A. et al. Star: ultrafast universal RNA-seq aligner. Bioinformatics 29, 15–21 (2013).

[107] Liao, Y., Smyth, G. K. & Shi, W. featurecounts: an efficient general purpose program for assigning sequence reads to genomic features. Bioinformatics 30, 923–930 (2014).

[108] Robinson, M. D., McCarthy, D. J. & Smyth, G. K. edger: a bioconductor package for differential expression analysis of digital gene expression data. Bioinformatics 26, 139–140 (2010).

[109] Ritchie, M. E. et al. limma powers differential expression analyses for RNA-sequencing and microarray studies. Nucleic acids research 43, e47–e47 (2015).

[110] Law, C. W., Chen, Y., Shi, W. & Smyth, G. K. voom: Precision weights unlock linear model analysis tools for RNA-seq read counts. Genome biology 15, R29 (2014).

[111] Zhou, X., Carbonetto, P. & Stephens, M. Polygenic modeling with Bayesian sparse linear mixed models. PLOS Genetics 9, e1003264 (2013).

[112] Guan, Y. & Stephens, M. Bayesian variable selection regression for genome-wide association studies and other large-scale problems (2011).

